# Chaperones Skp and SurA dynamically expand unfolded outer membrane protein X and synergistically disassemble oligomeric aggregates

**DOI:** 10.1101/2021.02.23.432414

**Authors:** Neharika Chamachi, Andreas Hartmann, Mai Quynh Ma, Georg Krainer, Michael Schlierf

## Abstract

Periplasmic chaperones Skp and SurA are essential players in outer membrane protein (OMP) biogenesis. They prevent unfolded OMPs from misfolding during their passage through the periplasmic space and aid in the disassembly of OMP aggregates under cellular stress conditions. However, functionally important links between interaction mechanisms, structural dynamics, and energetics that underpin both Skp and SurA association with OMPs have remained largely unresolved. Here, using single-molecule fluorescence spectroscopy, we dissect the conformational dynamics and thermodynamics of Skp and SurA binding to unfolded OmpX, and explore their disaggregase activities. We show that both chaperones expand unfolded OmpX distinctly and induce microsecond chain reconfigurations in the client OMP structure. We further reveal that Skp and SurA bind their substrate in a fine-tuned thermodynamic process via enthalpy–entropy compensation. Finally, we observed synergistic activity of both chaperones in the disaggregation of oligomeric OmpX aggregates. Our findings provide an intimate view into the multi-faceted functionalities of Skp and SurA and the fine-tuned balance between conformational flexibility and underlying energetics in aiding chaperone action during OMP biogenesis.

## Introduction

Molecular chaperones are key cellular components that play fundamental roles in maintaining cellular proteostasis^1,2^. Essential activities of chaperones include the assistance of *de novo* protein folding, the stabilization of non-native proteins in folding competent or unfolded states, and the rescue of misfolded and aggregated proteins^3–5^. Chaperones are an integral part of a wide range of protein quality control systems and their activities are intimately coupled to the biogenesis networks that aid the structural and functional maturation of proteins from their site of cellular synthesis to their target cellular compartments.

One biogenesis network where chaperone activity is of particular relevance is the biogenesis network of outer membrane proteins (OMPs)^6,7^. OMPs are a diverse group of β-barrel membrane proteins found in the outer membrane of Gram-negative bacteria, mitochondria, and chloroplasts. They fulfil a plethora of functions in cell signaling, metabolism, and transport^8–10^, are indispensable to the survival of bacteria^10,11^, and constitute important virulence factors and drug targets^12–14^. The OMP biosynthesis pathway is highly complex and conserved across all kingdoms of life^15^, and involves the coordinated action of a multicomponent protein machinery that aids in overcoming the many hurdles that these proteins have to surmount on their way to their target outer membrane^4,7,16^.

In Gram-negative bacteria, OMPs are translated in the cytoplasm, from where they are translocated across the inner bacterial membrane via the Sec machinery to the periplasmic space^17,5^. Within this aqueous compartment, OMPs are escorted to the outer membrane in an unfolded state (denoted as the uOMP state) with the aid of various chaperones that maintain the largely insoluble and aggregation prone uOMP polypeptide chains in a protected, partially unfolded state^18,19^. At the outer membrane, uOMPs are transferred to the β-barrel assembly machinery (BAM), which facilitates their native folding and insertion into the membrane^20^. Noteworthy, the periplasm is devoid of any known source of energy-providing molecules, such as ATP; hence, all chaperones as well as the entire folding machinery likely operates without the aid of external energy, following thermodynamic principles^21^.

Two chaperones which have been shown to be indispensable for the biogenesis of bacterial OMPs are the seventeen-kilodalton protein (Skp)^22^ and survival factor A (SurA)^23^. Skp and SurA, both located in the periplasm, exhibit anti-folding activity (also known as holdase activity), whereby they sequester uOMP substrates to prevent aggregation until they reach the bacterial outer membrane^24–26^. Their interaction with uOMPs is thermodynamically modulated due to the lack of energy-carrying molecules in the periplasm^27–29^. Depletion studies of periplasmic chaperones identified SurA as an essential chaperone for OMP biogenesis, leading to a drastic decrease in OMP density in the outer membrane due to the loss of SurA^30^. Skp depletion, on the other hand, led to an accumulation of misfolded OMPs and the activation of the cellular stress response^30^, while OMP density in the outer membrane remained the same. Interestingly, recent studies suggest substrate selectivity amongst the two chaperones^31^. Hence, it is of importance to understand the functional mechanisms underlying both Skp and SurA association with uOMPs.

Structural studies of the eight β-stranded protein outer membrane protein X (OmpX) in the presence of Skp using nuclear magnetic resonance (NMR) spectroscopy have found that unfolded OmpX (uOmpX) shows sub-millisecond backbone dynamics^32^ in complex with Skp. For binding of SurA to unfolded outer membrane protein A (uOmpA), both fluid globular^32^ and expanded states^31^ have been proposed. Recent studies have located various interaction sites of SurA and uOmpX using cross-linking, suggesting that SurA-bound uOmpX populates multiple conformations^31,33^. Yet, long-range polypeptide chain dynamics and conformational heterogeneity of unfolded OMPs upon binding to chaperones remain elusive. In particular, how SurA- and Skp-bound OMP dynamics and heterogeneities differ, given their differential roles in regulating protein folding in the periplasmic space. Dynamic aspects are hypothesized to be important for chaperone– uOMP interactions, particularly to fine-tune energetics of the binding reaction through a reduction of the entropic costs upon binding^28,29,34–37^. Yet, the enthalpic and entropic changes that determine Skp–OMP or SurA–OMP interactions and affect the conformations of the denatured substrate proteins are largely undefined.

In addition to the well-described holdase activities of Skp and SurA that protect OMPs from misfolding or aggregating, a recent study has suggested that Skp disaggregates oligomeric uOMP structures^38^. While SurA has not been directly implicated as a disaggregase, modelling studies propose a synergistic interaction amongst these chaperones especially under conditions of stress^18,39^, thus raising the question of the role of both chaperones played in disassembling OMP aggregates.

To gain insights into the multi-faceted functionalities of Skp and SurA and their action mechanisms, we study here the conformational dynamics and thermodynamics of the eight β-stranded protein OmpX in the presence of the chaperones, and explore their disaggregation activities. Using single-molecule Förster resonance energy transfer (smFRET), we resolve the heterogeneities, structural dynamics, and thermodynamics underlying the different states of uOmpX at near-native conditions. Strikingly, we find that both chaperones expand the unfolded polypeptide chain upon binding. The degree of expansion is concentration dependent for SurA, but not for Skp. Probing structural changes and chaperone interaction at different temperatures, we gain insights into the enthalpic and entropic contributions of complex formation and find that the interaction modes of both chaperones differ strongly and are dictated by entropy–enthalpy compensation. Finally, we use fluorescence correlation spectroscopy (FCS) to probe the disaggregation capabilities of Skp and SurA and find synergistic activity of both chaperones in the disassembly reaction of oligomeric OmpX aggregates. Our findings provide fundamentally new insights into the structural and energetic mechanisms underlying Skp and SurA chaperone–OMP interactions and their role in OMP biogenesis.

## Results

### Studying chaperone effects on uOmpX conformational dynamics under native-like conditions by smFRET

Probing chaperone effects and conformational dynamics of uOMPs under native-like conditions (i.e., in the absence of or at very low denaturant concentrations) has been difficult to achieve due to the aggregation prone nature of OMPs in aqueous environments^7,40,41^. Single-molecule methods, in particular single-molecule fluorescence techniques such as smFRET, operate at very low concentrations (i.e., in the pM range), and thus alleviate protein aggregation and precipitation challenges^42–44^. smFRET measurements further provide access to subpopulation-specific conformational heterogeneity, even at very low denaturant concentrations or the absence thereof^45^, and allow probing of long-range polypeptide chain dynamics across a wide spectrum of timescales in the presence of binding partners and without the need of synchronizing the ensemble with perturbation/relaxation techniques^46–48^. We therefore set out to develop an smFRET assay that allowed us to study the conformational dynamics of single uOmpX polypeptide chains in native-like aqueous environments and the effect that the chaperones Skp and SurA have on the structural and dynamic properties of uOmpX.

To begin, we prepared an N- and C-terminal fluorescently labelled double-cysteine OmpX variant (OmpX_1,149_) furnished with donor and acceptor dyes (Fig. 1a and Methods). The placement of the FRET-dye pair at the terminal ends of the protein allowed us to monitor the conformational properties of the uOmpX polypeptide chain. After preparation, labeling, and refolding of the protein in presence of the surfactant lauryldimethylamin-*N*-oxide (LDAO) (Supplementary Results, Supplementary Fig. 1a–c, and Methods), we subjected OmpX_1,149_ to a multi-step denaturation and dilution protocol to transfer it to a micelle-free aqueous buffer environment at pM concentrations (Fig. 1a). Briefly, we first denatured refolded OmpX_1,149_ with 6 M guanidinium chloride (GdmCl) at μM concentrations and subsequently diluted it in the presence of 6 M GdmCl to a concentration of 20 nM. This was followed by a thousand-fold dilution step into GdmCl-free buffer or into GdmCl-free buffer supplemented with the chaperones Skp or SurA. The resultant [GdmCl] after the final dilution step was 6 mM and the remaining [LDAO] was <100 nM, thus yielding uOmpX at 10 pM in a micelle-free native-like aqueous buffer that is largely devoid of denaturant (denoted as uOmpX_aq_). smFRET measurements were performed directly after the dilution protocol, thus mimicking the scenario when newly secreted OMPs enter the periplasm and encounter their interacting chaperones. Subsequently, fluorescence bursts were recorded from a large number of individual protein molecules diffusing through the confocal detection volume to generate FRET efficiency (*E*) histograms (see Methods). Because smFRET provides intramolecular distance information in the nanometer range by measuring the energy transfer between fluorescent donor and acceptor dyes attached to the polypeptide chain, *E* histograms report on the diversity of conformations with different levels of compactness (i.e., distance between the two dyes) and provide information on intrachain dynamics^44,49^.

**Figure 1.**
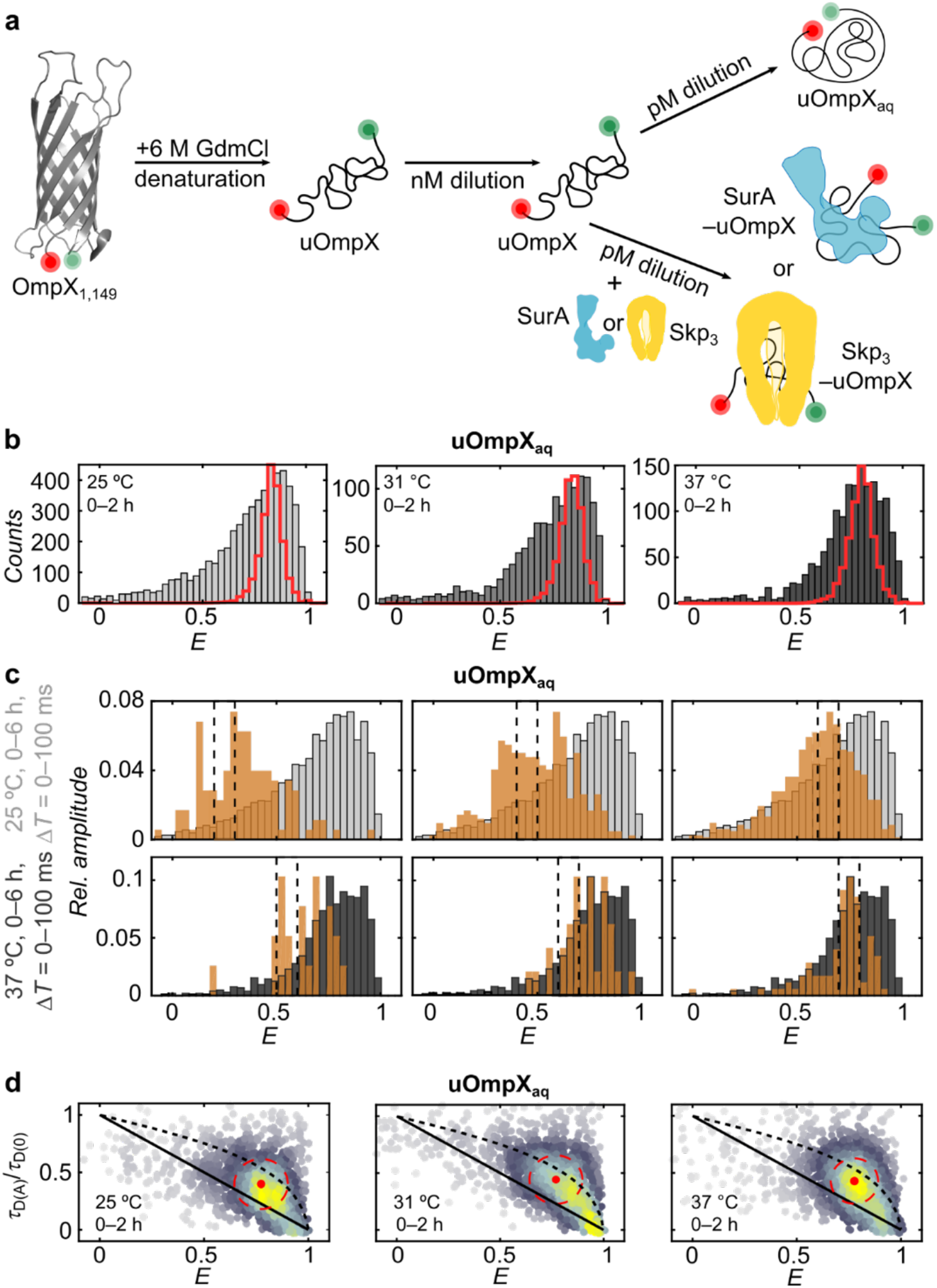
Probing structural dynamics of uOmpX under native-like conditions by smFRET. **(a)** Dilution scheme used for smFRET experiments to study unfolded and chaperone-bound uOmpX. The positions of donor (green sphere) and acceptor (red sphere) fluorophores at the N- and C-terminal ends of OmpX_1,149_ are indicated. **(b)** FRET efficiency (*E*) histograms of uOmpX in aqueous buffer conditions without denaturant (uOmpX_aq_) obtained during the first two hours of measurement at 25 °C, 31 °C, and 37 °C. Shot noise limited distributions are shown as red cityscapes. **(c)** Recurrence analysis with recurrence time intervals Δ*T* = {0, 100} ms and varying initial Δ*E* windows of 0.2–0.3, 0.4–0.5, and 0.6–0.7 at 25 °C (upper panel, black dashed lines) and 0.5–0.6, 0.6–0.7, and 0.7–0.8 at 37 °C (lower panel, black dashed lines). Recurrence histograms are shown in orange. Complete FRET efficiency histograms created from all detected bursts are shown in grey. **(d)** 2D scatter plot of relative fluorescence lifetime (*τ*_D(A)_/*τ*_D(0)_) vs *E* for uOmpX_aq_ at 25 °C, 31 °C, and 37 °C for 0–2 h of measurement. Lines represent the static FRET line (solid black line) and the expected correlation for a Gaussian chain (dashed black line). The red dot and the red dashed circle denote the center position and 68% area of the uOmpX_aq_ population. The scatter plot density is color coded (grey to yellow).

### uOmpX in the absence of chaperones is structurally heterogeneous and exhibits multi-tier dynamics

Under native-like buffer conditions and in the absence of chaperones, the *E* histogram of uOmpX_aq_ at 25 °C in the first two hours of measurement exhibited a broad distribution with a peak located at *Ê*_aq_ ≈ 0.77 and a tail spreading towards low transfer efficiencies (Fig. 1b, left panel). Broadened *E* distributions are generally ascribed to conformational heterogeneity with interconversion dynamics on timescales comparable to or longer than the observation time (~1–2 ms). Indeed, the width of the unfolded state population is in excess of that expected for a distribution that is limited only by detection shot noise (red cityscape in Fig. 1b), indicating the existence of a heterogeneous ensemble of conformations. Even at elevated temperatures of 31 °C and 37 °C (Fig. 1b, center and right panels, respectively), we observed a broadened *E* distribution further indicative of a rough energy landscape of the uOmpX_aq_ state^34^. Additionally, recurrence analysis^50^ revealed slow interconversions among the various sub-states on timescales longer than the burst duration (≥100 ms, see Fig. 1c). This suggests collective and transient conformational changes of remaining secondary motifs or tertiary long-range interactions, as have been described before for other OMPs under denaturing conditions^34,51^. Fluorescence lifetime and sufficient rotational averaging allowed us to exclude the possibility that the broadening originates from changes in dye quantum yields or a restricted rotational freedom of the dyes (see Supplementary Table 1). We also observed a modest expansion of the uOmpX_aq_ chain on the timescale of 4–6 hours (see Supplementary Results, Supplementary Fig. 1d–h, Supplementary Table 2), in line with reports which suggest that this mechanism prevents aggregation in the absence of both a membrane mimetic environment and chaperones^40^. Thus, our results indicate that uOmpX_aq_ might possess also an intrinsic non-aggregating ability.

In addition to the slow interconversion dynamics that give rise to broadened uOmpX_aq_ distributions, the unfolded polypeptide chain exhibited fast sub-millisecond reconfiguration dynamics at early time points (0–2 h), as indicated by a displacement of the uOmpX_aq_ population from the static FRET line in the relative donor lifetime (*τ*_D(A)_/*τ*_D(0)_, i.e., the ratio of the fluorescence lifetime of the donor in the presence and absence of acceptor, respectively) versus *E* plot (Fig. 1d). The uOmpX_aq_ population, however, did not fall onto a line that describes the intrachain distance distribution of a Gaussian chain, used to model completely unfolded polypeptides^52^ (Fig. 1d, dashed line). This substantiates our recurrence analysis^50^, in that, uOmpX_aq_ does not behave like a fully unfolded, random coil polypeptide. uOmpX_aq_ thus exhibits a multi-tier dynamical character with fast and slow intrachain motions, as has been previously observed for other proteins and membrane proteins^53–55^.

Interestingly, for the measurement conducted at 25 °C, at late time points (4–6 h), the *E* population of uOmpX_aq_, coincided with the static FRET line (Supplementary Fig. 1i, left panel), indicating that uOmpX_aq_’s fast reconfiguration dynamics vanish over time. This points toward the formation of long-range interactions in the unfolded-state and a ‘de-dynamization’ of the uOmpX_aq_ polypeptide chain while preserving the broad conformational heterogeneity. Such a behavior was absent in case of the measurement performed at 37 °C (Supplementary Fig. 1i, right panel), likely due to temperature-induced chain dynamics. Of note, in all experiments an additional minor population (~10%) at a high FRET efficiency (*E* ≈ 0.92) was observed. This very compact state is particularly observed after long measurement times of 4–6 h (Supplementary Fig. 1i) and likely constitutes a misfolded state, denoted as uOmpX_compact_ throughout the article.

In conclusion, we find that uOmpX_aq_ adopts a broad, heterogeneous ensemble of structures and populates a rugged energy landscape leading to a multi-tier dynamical behavior with both fast sub-millisecond chain reconfiguration dynamics and slow conformational interconversion. An overall expansion of the uOmpX_aq_ polypeptide chain is observed, in line with a conformational loosening mechanism that likely serves to alleviate uOMP aggregation^40^.

### Skp- and SurA-bound uOmpX is dynamically expanded

Next, we probed the effects that the chaperones Skp and SurA have on the structural and dynamic properties of the uOmpX polypeptide chain. To this end, we subjected denatured OmpX to a multi-step dilution and denaturation protocol to transfer the protein into native-like buffer in the presence of the chaperones Skp or SurA, and subsequently recorded *E* histograms (Fig. 1a).

In a first set of experiments, we complexed uOmpX with Skp at 37°C, which binds unfolded OMPs as a trimer^56,57^, denoted thereafter as Skp_3_. As reference, we first performed a measurement in the absence of the chaperone, and again observed a broad distribution at intermediate *E* values, representing the heterogeneous unfolded conformational ensemble of uOmpX_aq_, and a minor compact population at *E* values, representing uOmpX_compact_ (Fig. 2a; c.f. Fig. 1b). Upon addition of Skp_3_ to uOmpX, strikingly, an additional population at lower FRET efficiencies (*Ê ≈* 0.4) gradually arose, being already visible at 0.5 nM [Skp_3_] (Fig. 2b). At [Skp_3_] *≈* 2.5 μM, which is close to the reported periplasmic concentration of Skp^18,58,59^, the FRET efficiency distribution was dominated by this low *E* peak. The emergence of a third population at low FRET efficiencies, in addition to the two states of uOmpX in the absence of Skp_3_ (c.f. Fig. 2a), suggests that Skp_3_ interacts with uOmpX and forms a chaperone-bound state, in the following denoted as Skp_3_–uOmpX. This state is in equilibrium with and appears structurally distinct and more expanded than the uOmpX_aq_ and uOmpX_compact_ states.

**Figure 2.**
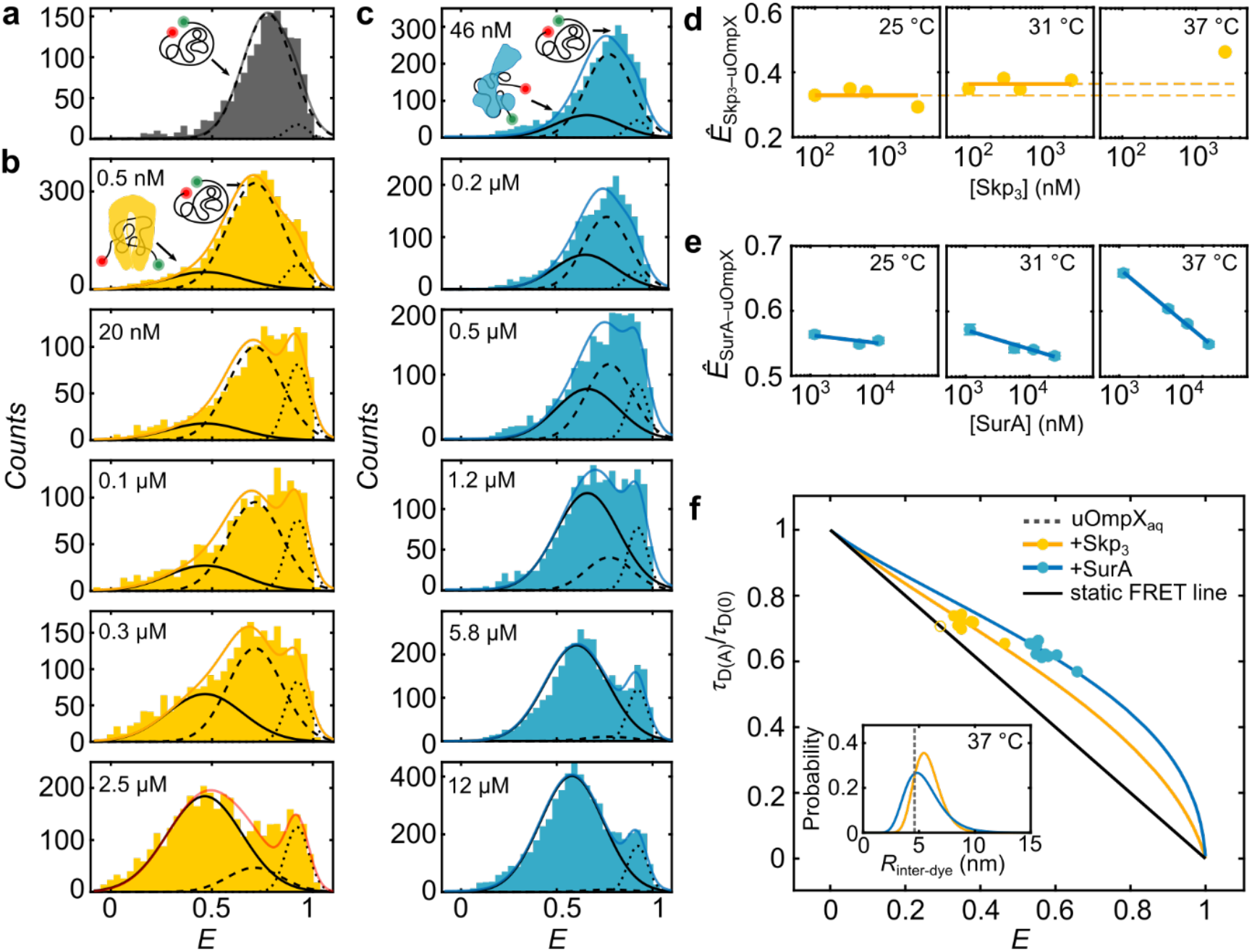
Conformations and structural dynamics of Skp- and SurA-bound uOmpX. **(a)** FRET efficiency (*E*) histogram of uOmpX in the absence of chaperones. **(b, c)***E* histograms of uOmpX in the presence of different concentrations of Skp_3_ (yellow) and SurA (blue), respectively. Concentrations are indicated. The three underlying Gaussian distributions are highlighted by black lines: Skp_3_- or SurA-bound uOmpX (Skp_3_–uOmpX or SurA–uOmpX, solid line), unbound uOmpX (uOmpX_aq_, dashed line) and compact uOmpX (uOmpX_compact_, dotted line). The sum of the three Gaussian distributions is shown in red. **(d)** FRET state peak positions (*Ê*) of Skp_3_–uOmpX and SurA–uOmpX at three different temperatures for measurements performed with [Skp_3_] > 100 nM and [SurA] > 1160 nM, respectively. **(e)** Relative fluorescence lifetime (*τ*_D(A)_/*τ*_D(0)_) and *E* position of Skp_3_–uOmpX and SurA–uOmpX (yellow and blue, respectively) populations for different temperatures. Here, we modelled the inter-dye distance with a log-normal distribution. The black line depicts the static FRET line. The inset shows the average inter-dye distance ⟨*R*_inter-dye_⟩ of uOmpX_aq_ (grey dashed line) and the empirical log-normal inter-dye distance distributions of uOmpX complexed with 2.5 μM Skp_3_ (yellow curve) and 11.6 μM SurA (blue curve), respectively.

We used Gaussian fitting to determine the three peak positions observed in the *E* histogram series (see Supplementary Fig. 2 and Methods). This analysis showed that the peak positions of the two populations in the high FRET regime exhibited *E* values of *Ê* ≈ 0.75 and *Ê* ≈ 0.92, which are in very good agreement with the *E* values of the uOmpX_aq_ (*Ê*_aq_ ≈ 0.77) and uOmpX_compact_ (*Ê_compact_* ≈ 0.92) states. This suggests that the presence of Skp_3_ does not affect the conformational properties of these states and implies that these do not directly interact with Skp_3_. We therefore identify these states, even in presence of chaperones, as uOmpX_aq_ and uOmpX_compact_. The FRET efficiency peak of Skp_3_-bound uOmpX was centered around *Ê*_Skp3–uOmpX_ ≈ 0.45, a considerably larger inter-fluorophore distance than the unbound uOmpX (i.e., uOmpX_aq_) state. A careful analysis of the donor lifetime, the fluorophore anisotropies, and the rotational correlation times of the fluorophores reassured that the lower *E* values reflect a global conformational change arising from the expansion of the uOmpX polypeptide chain upon binding to Skp_3_ (Supplementary Table 1). Moreover, Gaussian fitting revealed that, at [Skp_3_] = 2.5 μM, ~80% of uOmpX molecules were in a Skp_3_–uOmpX complexed state, suggesting that the presence of Skp shifts the equilibrium from the unbound to the bound form.

In a second set of experiments, we diluted uOmpX in the presence of the chaperone SurA at 37°C. Upon addition of SurA, we observed, similarly to Skp_3_, an additional FRET efficiency peak at lower *E* values (Fig. 2c). Increasing [SurA] from 46 nM to 25 μM, lead to an increase of the low *E* state population (i.e., SurA–uOmpX). Gaussian fitting allowed us to identify the peak position and fraction of SurA-bound uOmpX with *Ê*_SurA–uOmpX_ ≈ 0.6 at a concentration of ~6 μM, which is close to the intracellular concentration of SurA^18,59^. At this concentration, ~87% of uOmpX molecules are bound to the chaperone. The lower *Ê* value of SurA–uOmpX indicated also an expanded conformation as compared to uOmpX_aq_, yet a less expanded conformation as compared to Skp_3_–SurA. The distinct peak positions of the two chaperone-bound states (*Ê*_Skp3–uOmpX_ ≈ 0.45 and *Ê*_SurA–uOmpX_ ≈ 0.6) thus suggest that they themselves occupy a distinguishable configurational space likely due to the difference in the interaction mechanisms between the chaperone and its substrate^7,16^.

Since temperature affected the conformation and reconfiguration dynamics of uOmpX_aq_, we asked how temperature influences the structural ensemble and dynamics of chaperone-complexed uOmpX. To this end, we performed titration experiments at 31 °C and 25 °C and evaluated *Ê*_Skp3–uOmpX_ and *Ê*_SurA–uOmpX_ at different [Skp_3_] and [SurA] (Supplementary Fig. 3 and Supplementary Table 3). The FRET efficiencies of the unbound (uOmpX_aq_) and compact uOmpX (uOmpX_compact_) states did not vary markedly across the chaperone concentration series performed at different temperatures (Supplementary Fig. 4). For the chaperone-bound fraction of uOmpX, *Ê*_Skp3–uOmpX_ remained constant, within error, across nearly three orders of magnitude of [Skp_3_] at 25 °C (Fig. 2d, left panel). This suggests a concentration-independent, possibly 1:1 stoichiometry of Skp_3_ and uOmpX, in accordance with previous studies^32,60^. Increasing the temperature of the complex, however, shifted *Ê*_Skp3–uOmpX_ to higher *E* values (Fig. 2d), while at each temperature, *Ê*_Skp3–uOmpX_ itself remained constant again across all concentrations. We infer that, at elevated temperatures, uOmpX in complex with the chaperones appears to form a more compact structural ensemble, possibly due to increased dynamics of uOmpX in the Skp_3_ cavity at 37 °C as compared to 25 °C, similar to recent observations made by NMR spectroscopy^32^. By contrast, for SurA–uOmpX complexes, *Ê*_SurA–uOmpX_ decreased with increasing chaperone concentrations at 25 °C (Fig. 2e, Supplementary Table 4). Interestingly, at elevated temperatures, the dependency on [SurA] was enhanced and we observed a strong decrease of *Ê*_SurA–uOmpX_ with increasing [SurA] at 37 °C from *Ê*_SurA–uOmpX_ (1.2 μM) ≈ 0.66 to *Ê*_SurA– uOmpX_ (25 μM) ≈ 0.55. This suggests that uOmpX is more expanded at elevated [SurA], which can be explained by either a different interaction mode or that uOmpX is sequestered by multiple SurAs as also suggested by previous studies^31,33^.

Next, we modeled the structural ensemble of the states globally using a log-normal distribution and obtained a standard deviation of the donor–acceptor distance (*σ_R_*) and the expected inter-dye distance ⟨*R*_inter-dye_⟩ (inset Fig. 2f; Methods). A measure of heterogeneity among a population of varying end-to-end distance is the coefficient of variance, 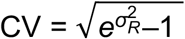, where a larger CV indicates an increased heterogeneity. We determined CVs of 0.204 ± 0.036 and 0.304 ± 0.022 for Skp_3_-bound and SurA-bound uOmpX, respectively. Compared to SurA–uOmpX, the CV of the Skp_3_–uOmpX complex is reduced and indicates less conformational heterogeneity for the bound substrate, likely due to the binding of uOmpX in the cavity of Skp_3_^56,57^. By contrast, the higher CV for the SurA–uOmpX complex marks an increased heterogeneity. This increase in heterogeneity could originate from the transient multi-site binding of SurA resulting in diverse unstructured and expanded uOmpX conformations with none, one, or even multiple SurAs bound. We further used the log-normal distribution to extract inter-dye distances in order to quantify the expansion of the chaperone-bound uOMP complexes. Compared to uOmpX_aq_, the inter-dye distance grows from 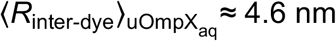 to 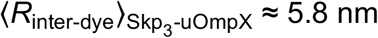 at 2.5 μM [Skp_3_] and to 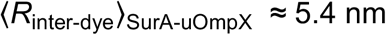 at 12 μM [SurA].

In a last step, to gain further insights into the structural and dynamical properties of chaperone-bound uOmpX, we analyzed the underlying dynamics of the chaperone-complexed uOmpX polypeptide chain. To this end, we extracted single-molecule events of the chaperone-bound population using a combined FRET efficiency and relative donor fluorescence lifetime filter (Supplementary Fig. 2). We determined the peak positions for *E* and *τ*_D(A)_/*τ*_D(0)_ for each condition and plotted the values in a 2D diagram (Fig. 2f and Supplementary Table 2). The bound FRET efficiencies (*Ê*_Skp3–uOmpX_ and *Ê*_SurA–uOmpX_) across different chaperone concentrations were offset in distinct clusters from the static FRET line. The offset indicates that the polypeptide chain undergoes fast configurational changes on the sub-millisecond timescale while being bound to chaperones.

To explore the timescale of dynamics further, we implemented species-filtered 2D fluorescence lifetime correlation spectroscopy (2D FLCS)^61,62^ and extracted characteristic chain reconfiguration times. Briefly, in a first step, photon pairs of FRET efficiency-filtered fluorescence bursts with a time gap matching the time interval of Δ*T* and a window size of 2 × ΔΔ*T* (Fig. 3a) were sorted according to the microtime of the initial and final photon (*t*_1_ and *t*_2_, respectively) in the two-dimensional emission-delay histogram (Supplementary Fig. 5). The timescale of interconversion dynamics between different molecular states were then extracted by comparing slices of the 2D histogram with either short or long initial microtimes (blue and red, respectively, in the center panel of Fig. 3a and in the histograms of Fig. 3b, c and d). If the chosen Δ*T* matches the time of a conformational change, the emission-delay histograms for two initial microtime windows will start to separate. For uOmpX_aq_, all delay time windows Δ*T ±* ΔΔ*T* produced separated emission-delay histograms (Fig. 3b). The amplitude of separation was greater at a lower time point Δ*T* = 5 μs as compared to that at higher time points of 5 μs or 240 μs. This reinforced our finding that uOmpX_aq_ exhibits both fast and slow conformational dynamics by virtue of its rough energy landscape. For both Skp_3_–uOmpX (Fig. 3c) and SurA–uOmpX (Fig. 3d) complexes, by contrast, the emission-delay histograms started separating at lower time points Δ*T* = 2.5 μs and Δ*T* = 6 μs, respectively. Skp- or SurA-bound uOmpX thus shows a dominating microsecond conformational change, suggesting that both chaperones modulate the energy landscape of uOmpX such that the energy barriers between the sub-population of their bound substrate are reduced, leading to a dynamization of the polypeptide chain in the chaperone-bound state.

**Figure 3.**
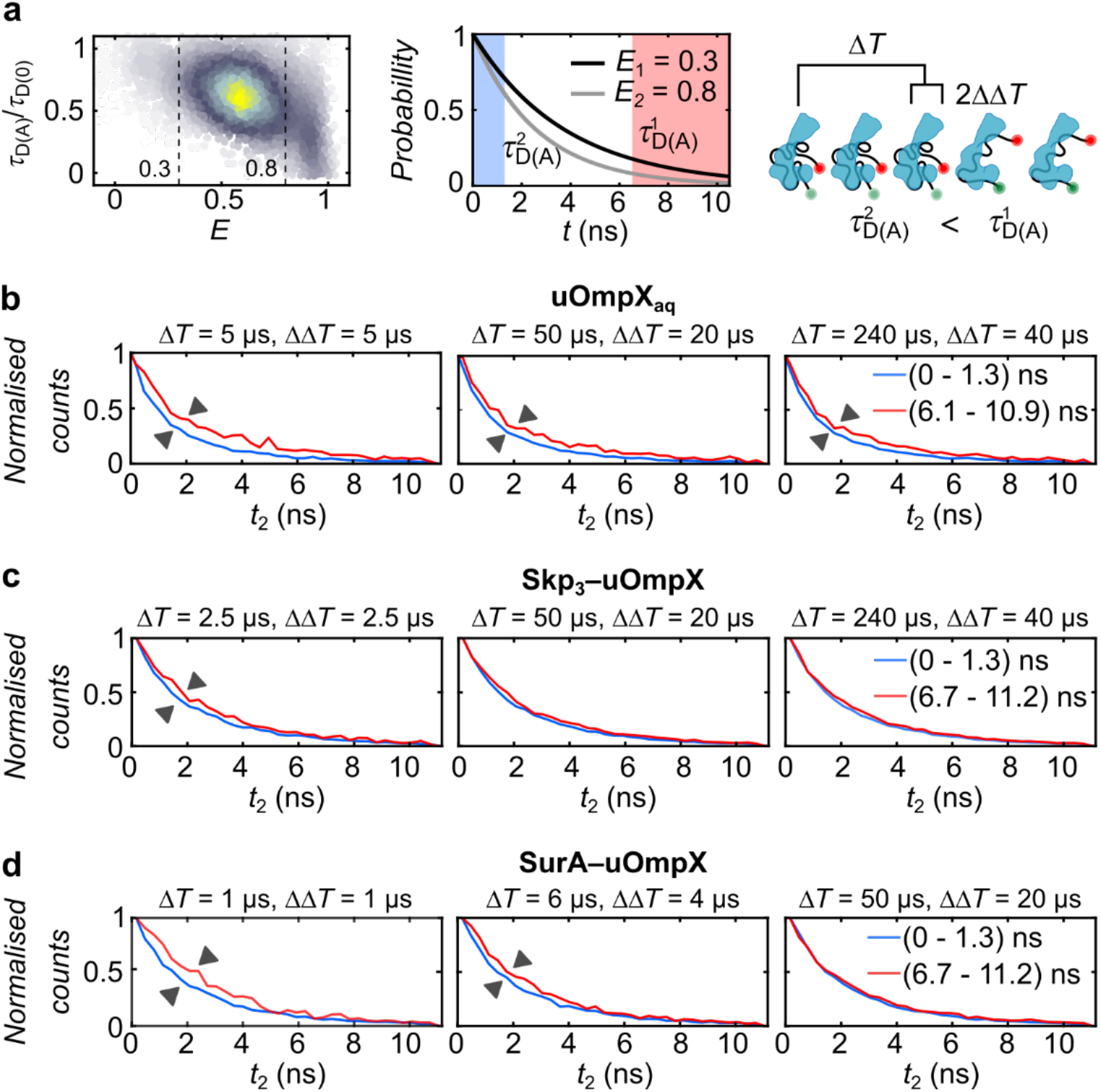
Conformational dynamics of the Skp_3_- and SurA-bound uOmpX. **(a)** Representation of species-filtered 2D FLCS using SurA–uOmpX complexes as an example. Here, *t* is the microtime of the emitted photons, Δ*T* is the time interval and ΔΔ*T* is the window size. Emission-delay histograms corresponding to **(b)** uOmpX_aq_, **(c)** Skp_3_-uOmpX and **(d)** SurA-uOmpX. The legend indicates the short and long initial microtimes.

To conclude, we found that binding of the chaperones Skp_3_ and SurA leads to an expansion of the uOmpX polypeptide chain. Skp_3_-complexed uOmpX showed a decreased conformational heterogeneity, while binding of SurA increased the conformational heterogeneity of uOmpX, indicating very distinct interaction mechanisms of the chaperones on their substrates. Complexed uOmpX exhibited fast chain reconfiguration dynamics on timescales < 10 μs, unlike uOmpX_aq_ alone, which showed multi-tier dynamics on a large range of timescales (sub-millisecond to ≥100 ms) originating from its rugged energy landscape. The fast dynamics and the expansion of chaperone-bound uOmpX may aid in preventing uOmpX from getting trapped in a fortuitous conformation which might be folding-incompetent or inaccessible to the BAM complex. A wealth of different conformations also indicates a small entropic penalty for complexing uOmpX to Skp_3_ or SurA.

### Entropic and enthalpic contributions of chaperone–uOmpX interactions

We next set out to obtain insights into the energetic contributions of Skp and SurA chaperone–uOmpX interactions. Interaction isotherms at different temperatures allow for the extraction of enthalpic, Δ*H*, and entropic, Δ*S,* contributions to the free energy change of interaction, Δ*G,* which are connected to the association constant, *K*_a_, for a bimolecular association reaction via:

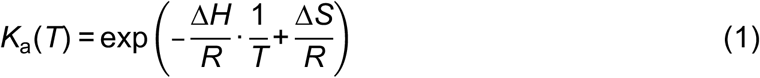

In order to extract entropic and enthalpic contributions, we developed a global analysis approach that allows reconstruction of FRET efficiency distributions at different temperatures and chaperone concentrations (Fig. 4a and Methods). Briefly, by varying the optimization parameters enthalpy (Δ*H*), entropy (Δ*S*), and considering the fraction of the compact state (*f*_c_(*T*)), which is likely unable to be chaperone complexed, as well as the maximum fraction of the bound state (*f*_b,max_(*T*)), and minimizing the reduced chi-square, the best estimators for Δ*H* and Δ*S* were obtained. Here, we chose a Hill coefficient of 1 for Skp_3_ and 1.5 for SurA, based on earlier reports^32,33,60,63^. We further assumed that Skp_3_ remained trimeric under our experimental conditions (see Supplementary Information Methods). At 37 °C, we obtained a *c*_1/2_ =(359 ± 0.1) nM and (407 ± 0.1) nM for Skp_3_–uOmpX and SurA–uOmpX (Supplementary Table S4), respectively, in good agreement with previously reported values for the chaperone–OMP interactions^33,36,63^. More interestingly, for Skp_3_–uOmpX complex formation, the binding enthalpy and entropy changes were Δ*H* (Skp_3_–uOmpX) = –298 ± 19 kJ mol^−1^ and Δ*S* (Skp_3_–uOmpX) = –0.84 ± 0.06 kJ mol^−1^ K^−1^ (Fig. 4b), respectively, yielding an interaction free energy change of Δ*G* (Skp_3_–uOmpX) = (−38.3 ± 0.2) kJ mol^−1^ at 37 °C. The change of enthalpy upon binding is large, in agreement with recent findings demonstrating rich interaction of the uOmpX polypeptide chain with the cavity formed by Skp_3_^32^. At the same time, the change in interaction entropy is strongly negative, suggesting a stark reduction of the overall configurational space for uOmpX upon its encapsulation by Skp_3_. Hence, Skp_3_ binding to uOmpX is entropically unfavorable, yet this entropic penalty is counter-weighted by a large enthalpic contribution, enabling the binding of Skp_3_ to the client OMP.

**Figure 4.**
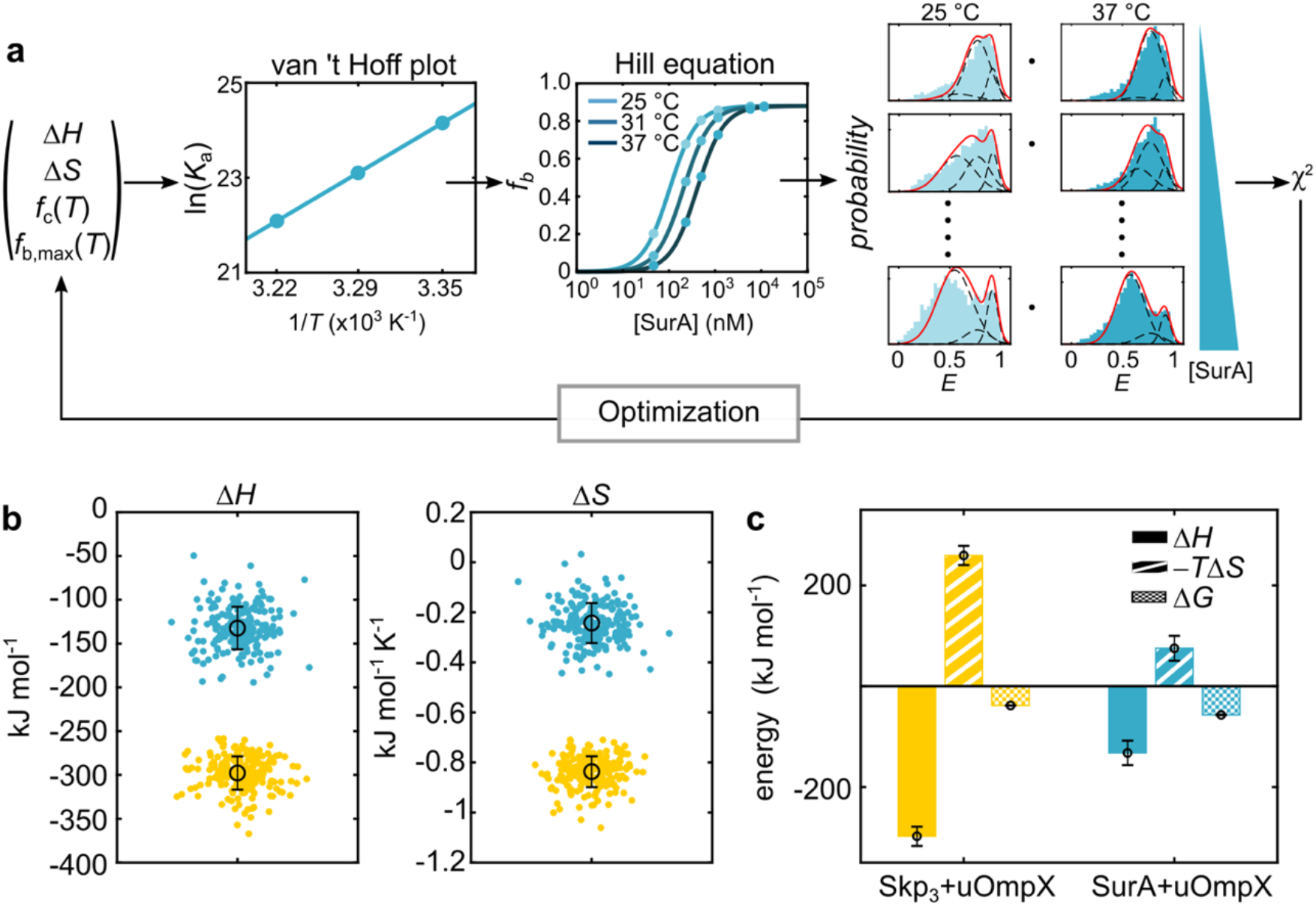
Thermodynamics of Skp_3_ and SurA interaction with uOmpX. **(a)** Schematic of the global chi-square minimization routine to obtain the enthalpic change (Δ*H*) and entropic change (Δ*S*) for Skp_3_ and SurA interaction with uOmpX. Δ*H*, Δ*S*, *f*_c_(*T*) (i.e., fraction of compact uOmpX at each temperature) and *f*_b,max_(*T*) (i.e., maximum fraction of Skp_3_–uOmpX or SurA–uOmpX) were the varying fitting parameters. ln(*K*a) is the natural logarithm of the association constant, *T* is the temperature in Kelvin (K) and [SurA] is the initial SurA concentration in nM. **(b)** Δ*H* and Δ*S* values for Skp_3_ and SurA interaction with uOmpX as obtained by a bootstrapping algorithm. The best estimators for Δ*H* and Δ*S* are indicated as a black circle with standard deviation. **(c)** Comparison of the best estimators for Δ*H, T*Δ*S*, and Δ*G* for Skp_3_ and SurA interaction with uOmpX. Δ*G* is the free energy change of interaction at 37 °C.

For SurA–uOmpX complex formation, the binding enthalpy and entropy changes were Δ*H* (SurA–uOmpX) = –132 ± 24 kJ mol^−1^ and Δ*S* (SurA–uOmpX) = –0.24 ± 0.08 kJ mol^−^ 1 K^−1^ (Fig. 4b), respectively, yielding an overall stabilization of the complex of Δ*G* (SurA– uOmpX) = (−57 ± 1) kJ mol^−1^ at 37 °C, which is slightly more favorable than for Skp_3_. The change of enthalpy was two-fold lower than for Skp_3_, likely due to more transient binding as suggested in recent studies^31,33^. At the same time, the change of entropy of uOmpX upon SurA binding was three-fold lower than with Skp_3_, and close to zero, indicating only a minimal change of configurational space for the binding of SurA to uOmpX. This agrees well with our finding of an increased coefficient of variation (i.e., structural heterogeneity) for SurA–uOmpX as compared to Skp_3_–uOmpX. Thus, binding of SurA to its client uOmpX is markedly different in nature than Skp_3_ binding to uOmpX. For Skp_3_, the reduction of conformational freedom of uOmpX in the cavity of Skp_3_ needs to be compensated by an increased interaction enthalpy (Fig. 4c). By contrast, uOmpX binding to the surface of SurA reduces the conformational freedom to a lesser extent and thus a lower enthalpic contribution is sufficient to enable binding (Fig. 4c).

### SurA and Skp_3_ act synergistically as disaggregases on oligomeric OmpX structures

Aside from the known uOMP holdase activities of Skp and SurA, recent work has suggested that Skp also has the ability to act as a disaggregase to disassemble oligomeric OMP structures^38^. By contrast, only indirect evidence^18,39^ suggests that SurA might be involved in synergetic interaction with other periplasmic chaperones, such as Skp, under cellular stress in the disassembly of aggregates.

To address this question, we investigated the action of Skp_3_ and SurA towards aggregated OmpX (OmpX_Agg_) and devised an FCS-based assay that allows probing the disaggregase efficacy of the two chaperones. To this extent, we diluted labeled and denatured uOmpX at pM concentrations with unlabeled and denatured uOmpX in buffer to yield a final OmpX concentration of 1 μM (Fig. 5a). The residual GdmCl and LDAO concentrations were 12 mM and <200 nM, respectively. It is important to note, that due to the 100,000-fold concentration difference of labeled and unlabeled uOmpX, only a small fraction of potential aggregates will be fluorescently marked and thus visible to FCS measurements. We incubated the mixture for 10 min and subsequently performed FCS measurements at 37 °C and obtained a multimodal auto-correlation function with two major components suggesting the presence of a fast and a slowly diffusing species (Fig. 5b, green). Comparison with an FCS curve recorded from a highly diluted sample of only labeled uOmpX_aq_ (~10 pM) indicates that the fast-diffusing component in the correlation curve of the aggregation sample stems from monomeric uOmpX_aq_, with a characteristic diffusion time of *τ*_Diff_ = (0.27 ± 0.01) ms (Fig. 5b, grey, Supplementary Table 5, Supplementary Fig. 6a). The second, slowly diffusing species in the OmpX_Agg_ measurements showed a *τ*_Diff_ = (31.6 ± 1.3) ms, about two orders of magnitude slower than single uOmpX_aq_ molecules. Notably, OmpX_Agg_ is possibly an ensemble of differently sized aggregates; hence, values reported here reflect average values. We determined that in our aggregation mixture, about 24 ± 2% of the labeled uOmpX molecules are part of aggregates (termed apparent fraction, Fig. 4d). Interestingly, at lower temperatures, the propensity of OmpX aggregation was reduced (Supplementary Fig. 6e).

**Figure 5.**
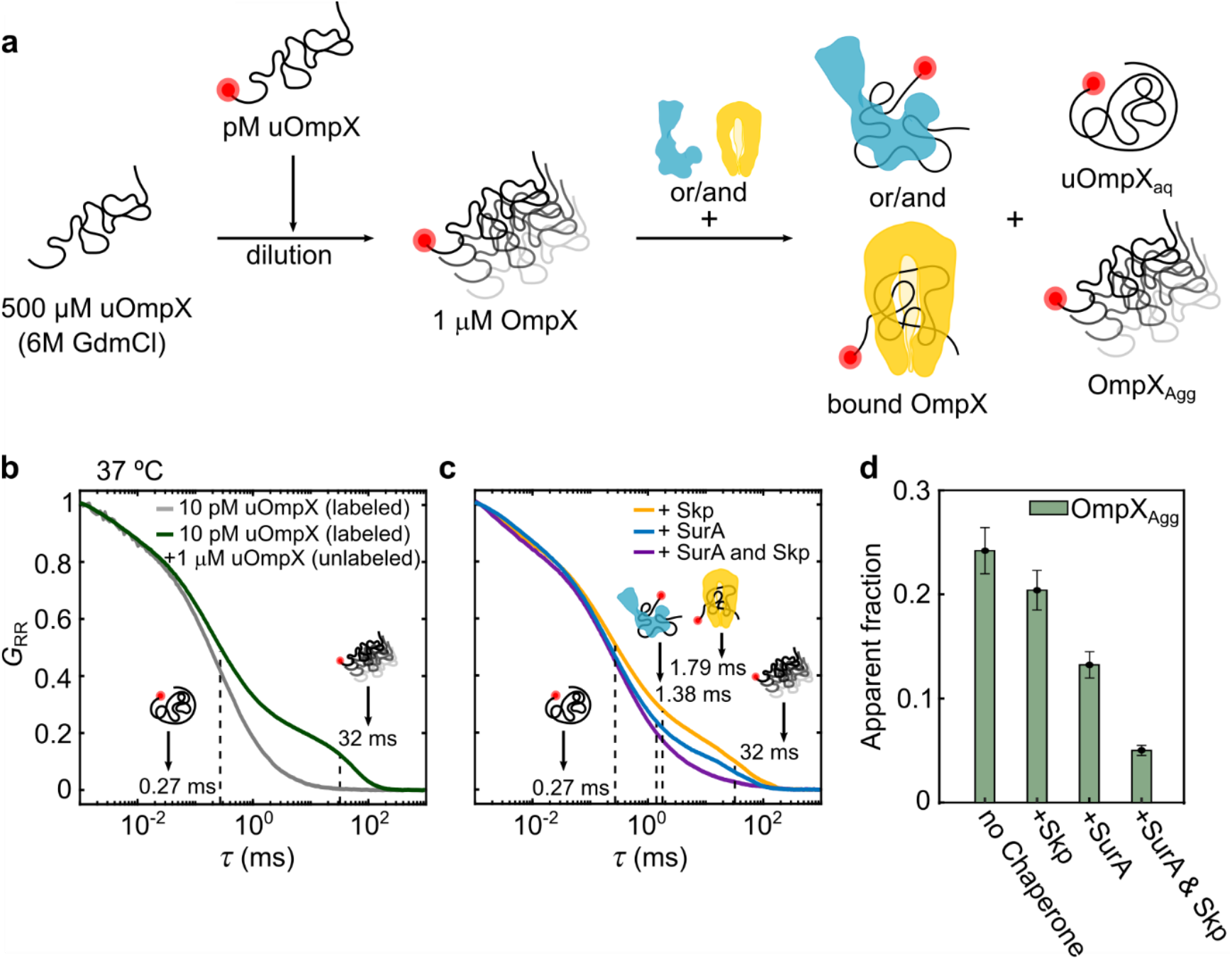
Disaggregation of OmpX aggregates by Skp and SurA. **(a)** Dilution and complex formation scheme to investigate OmpX disaggregation by Skp and SurA. **(b)** FCS curves corresponding to measurements containing 10 pM uOmpX (grey) and 1 μM uOmpX + 10 pM uOmpX (green) in aqueous buffer conditions without chaperones yield two species: uOmpX_aq_ and/or OmpX_Agg_ with diffusion times as indicated. Here, *G*RR is the normalized autocorrelation function of the acceptor dye and τis the diffusion time. **(c)** Upon addition of Skp (yellow) or SurA (blue) or both (purple) to the aggregate mixture, more species appear corresponding to Skp_3_–uOmpX and/or SurA–uOmpX complex with diffusion times as indicated. **(d)** The probability of OmpX_Agg_ (green bar) in different measurement conditions: at μM concentration of OmpX in aqueous buffer without chaperones, at μM concentration of OmpX in presence of Skp (+Skp), at μM concentration of OmpX in presence of SurA (+SurA) and at μM concentration of OmpX in presence of both the chaperones (+ SurA and Skp).

Next, we studied the effect that the chaperones Skp and SurA have on OmpX aggregation. To this end, we first incubated the mixture containing 1 μM unlabeled uOmpX and 10 pM labelled uOmpX for 10 min and then added either 2.5 μM Skp_3_ or 5.8 μM SurA to the mixture. We recorded FCS curves at 37 °C directly after the dilution and obtained multimodal curves, indicating the presence of more than two components in solution. In addition to the fast-diffusing species with *τ*_Diff_ = (0.27 ± 0.01) ms reflecting free uOmpX_aq_, and the aggregated OmpX_Agg_ species with very high diffusion times (*τ*_Diff_ ≈ (32 ± 1) ms), we expected to observe also chaperone-complexed uOmpX species with intermediate diffusion times. By comparison with an FCS curve recorded from a highly dilute sample of labeled uOmpX_aq_ (~10 pM), we assigned diffusion times of *τ*_Diff_ = (1.79 ± 0.05) ms and (1.38 ± 0.03) ms to Skp_3_- and SurA-bound monomeric uOmpX, respectively (Supplementary Fig. 6b and c). To quantify the fractional amounts of the three components (i.e., uOmpX_aq_, chaperone-bound monomeric uOmpX, and OmpX_Agg_), we fitted the autocorrelation data of our mixture experiment with three components. Here, the respective diffusion times were kept constant to extract the amplitudes of each component as they are inversely proportional to the concentrations and fractional amounts of the species. Addition of Skp_3_ to the aggregation mixture reduced the apparent fraction of OmpX_Agg_ to 20 ± 2 % and addition of SurA even to 13 ± 1 %. Interestingly, in both cases the disaggregation of OmpX was found to be accompanied by an increase in the free uOmpX_aq_ population (Supplementary Fig. 6f) and only negligibly in the chaperone–uOmpX complex. Assuming, uniform mixing of labelled and unlabeled uOmpX sample, the increase of the free uOmpX population suggests that the chaperones show a higher affinity towards OmpX_Agg_ as compared to uOmpX. This finding is significant, as it suggests that careful adaptation of affinities is crucial during physiological stress conditions. In such a scenario, disaggregation activity might be more important than holdase activity to prevent the formation of large toxic OMP aggregates in the periplasm.

Finally, we asked the question whether there are synergistic effects on the disaggregation reaction, given that both chaperones are present in the periplasm. We performed our disaggregation assay in which both chaperones were added simultaneously to the mixture containing OmpX_Agg_ and labelled uOmpX at pM concentrations after 10 min incubation at 37°C. Strikingly, the fraction of OmpX_Agg_ was reduced to ~5% (Fig. 5d). Taking into account our single-chaperone measurements, for an independent, additive reduction of the apparent OmpX_Agg_, we would have expected a reduction to ~9%. The increased disaggregation by a factor of two for both chaperones together suggests therefore a cooperative action of Skp_3_ and SurA.

## Conclusions

Periplasmic chaperones Skp and SurA are essential players in OMP biogenesis. They act as holdases, preventing uOMP substrates from misfolding, and exhibit disaggregase functionalities, aiding in the clearance of OMP aggregates^32,38^. Understanding the molecular action mechanisms of these chaperones is fundamental to unravelling the functional role that these chaperones play during OMP biogenesis. Here, we have provided an intimate view on the structural, dynamic, and thermodynamic aspects of uOmpX in complex with Skp and SurA, and explored their disaggregase activities.

Using smFRET, we have characterized the structural dynamics of chaperone-free and chaperone-bound uOmpX under near-native conditions and show that both chaperones, Skp and SurA, expand uOmpX upon binding. While previous reports have observed an expansion for SurA-bound OMPs^31,38^, an expansion upon Skp binding of an OMP in its cavity has not been observed. More surprisingly, the Skp_3_-bound uOmpX state is more expanded as compared to the SurA–uOmpX state over the concentration range tested, suggesting that uOmpX interacts with Skp_3_ at multiple sites by spreading across the inner surface of the chaperone cavity (Fig. 6a). The expansion of uOmpX upon Skp_3_ binding was independent of the [Skp_3_], indicating that the valency of Skp_3_ likely does not change as expected for the small eight β-stranded OmpX^60^. The SurA–uOmpX complex (Fig. 6b), by contrast, showed an increased expansion with increasing [SurA], which is in line with multiple SurAs binding to one uOmpX. The higher valency was previously already suggested with a Hill-coefficient of *n* = 1.5 for SurA–uOmpX association^33^. Interestingly, chaperone-induced expansion was also observed for the ATP-dependent DnaJ–DnaK system for cytosolic proteins as clients^64^, by forming a chain of DnaK molecules on the denatured substrate. However, unlike for Skp_3_ and SurA, for DnaK, the dynamics of interaction were largely dominated by ATP hydrolysis. Mechanistically, an expansion of OMPs by either Skp or SurA likely prevents intra-chain contacts and, by extension, misfolding of uOmpX in the aqueous periplasmic space. Additionally, chain expansion can aid in exposing the β-signal peptide sequence, which facilitates recognition by BAM^65,66^. Hence, our results point to a general mechanism of chain expansion, which in the case of Skp and SurA, enables OMP chains to defer misfolding even in the absence of a high energy substrate.

**Figure 6.**
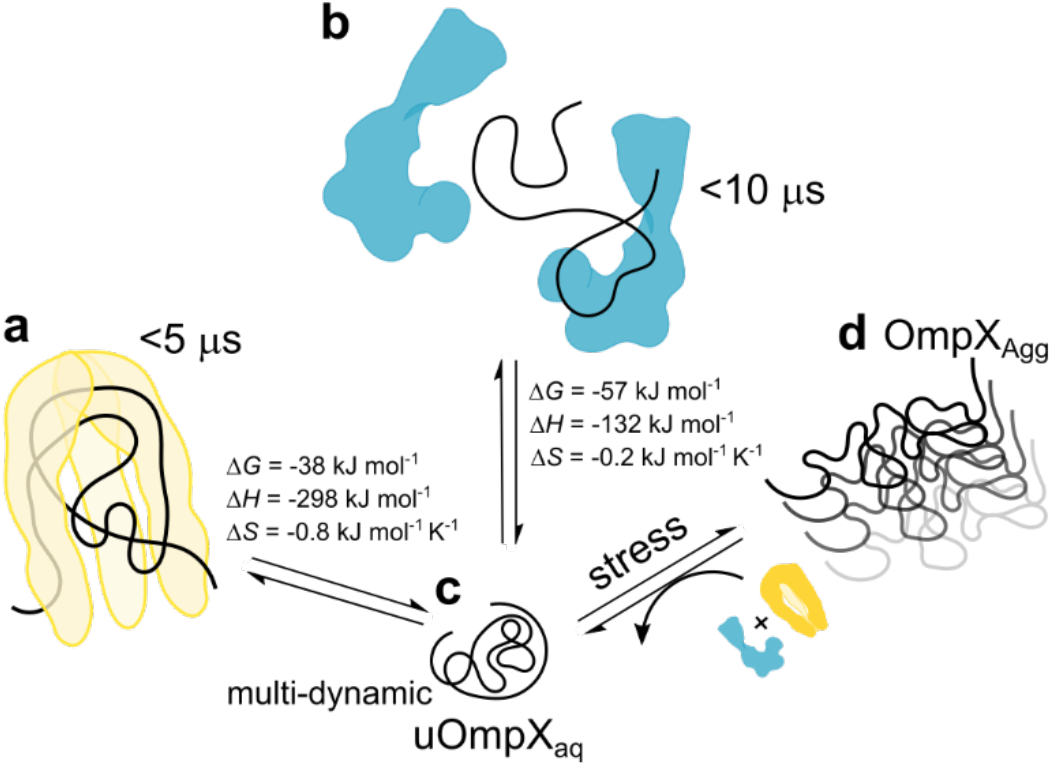
Model of Skp_3_ and SurA chaperone action on uOmpX and OmpX_Agg_. **(a)** Skp_3_–uOmpX is expanded due to numerous intermolecular interactions between the chaperone and its substrate, while the substrate itself undergoes fast chain reconfiguration on timescales <5 μs. **(b)** The increased expansion of uOmpX upon incrementing [SurA] indicates that more than one molecule of SurA binds to uOmpX. Similar to Skp_3_, SurA expands its substrate and induces uOmpX chain reconfiguration on a timescale of <10 μs. Interactions of both chaperones with uOmpX are energetically calibrated through an exquisite entropy– enthalpy compensation, as indicated by the values of entropy and enthalpy change (Δ*H* and Δ*S*, respectively), along with the free energy of interaction (Δ*G*). **(c)** uOmpX_aq_ exhibits both sub-millisecond chain reconfiguration dynamics and ≥100 ms timescale conformational changes in aqueous buffer. **(d)** Both chaperones can disassemble aggregated OmpX, which can emerge under stress conditions.

We further quantified the dynamics of both Skp_3_- and SurA-bound uOmpX using 2D-FLCS. We demonstrate that the intrachain dynamics of the chaperone-complexed uOmpX chain are on a timescale of 5 to 10 μs (Fig. 6a and 6b), whereby the fast chain reconfiguration dynamics are possibly facilitated through local transient interactions with the chaperones. This is in contrast to the multi-tier (i.e., both slow conformational and fast reconfiguration chain) dynamics observed for the aqueous uOmpX state (Fig. 6c). Periplasmic chaperones thus increase conformational flexibility, despite the expansion of the unfolded state. We hypothesize that such a manipulation and dynamization of the energy landscape of the OMP substrate might be beneficial for BAM interactions^65^. Notably, transient binding interactions and multi-valency interactions were also observed for the cytoplasmic chaperone Trigger Factor (TF), which is structurally homologous to SurA, and like the latter also involved in ATP-independent transfer of clients to the downstream chaperone machinery for folding^67^.

Given that the periplasm is deficient of ATP as an energy source, protein folding and chaperoning, involving binding to the client uOMPs, has to be driven entirely by a change in the free energy of the system. An analysis of binding isotherms at different temperatures revealed that the change of entropy upon binding to the client uOmpX substrate was low for Skp_3_ and close to zero for SurA, suggesting that the conformational freedom is affected only minimally. A low entropic change upon binding has been hypothesized to be favorable for locally transient chaperone–uOMP interactions, due to a reduced enthalpy–entropy compensation^29^. Strikingly, comparing Skp_3_ and SurA, we find that the chaperone-uOmpX interactions are fine-tuned exquisitely, meaning a higher entropic change of Skp_3_ is compensated by a higher enthalpic term, or vice versa in the case for SurA, to ensure a favorable, yet low change of free energy upon binding (Fig. 6). Hence, the different contributions of enthalpy and entropy to both complexes suggests that the thermodynamics of the Skp_3_–uOmpX and SurA–uOmpX interaction are differently calibrated in accordance with their chaperoning mechanisms.

Lastly, we found that both chaperones have the ability to disassemble aggregated OmpX (Fig. 6d). Interestingly, both chaperones show a higher affinity towards aggregated OmpX than uOmpX. A higher affinity towards the aggregate might be particularly useful when aggregated OmpX emerges, for example, under stress conditions to avoid the accumulation of OMP aggregates. Surprisingly, we found that Skp_3_ and SurA act synergistically on uOmpX aggregates, suggesting higher order interactions between both periplasmic chaperones. Disaggregation mechanisms have also been observed in the ATP-dependent cytoplasmic chaperones of Hsp70 family^68,69^. However, both Skp and SurA disaggregate oligomeric structures in the absence of an external energy source, thereby supporting the known disaggregase DegP, with a yet to be determined mechanism.

In summary, both Skp and SurA binding of uOmpX expands the unfolded polypeptide chain distinctly. Interestingly, the unfolded polypeptide chain, in complex with both chaperones, exhibits a dynamization on the microsecond timescale. While the structural features of the bound polypeptide appear similar, the thermodynamic aspects of the Skp_3_–uOmpX and SurA–uOmpX interaction differ and SurA binding results in a reduced loss of entropy compared to Skp binding. In a biological context, expansion and fast chain reconfiguration of the bound polypeptide may serve to ensure that the uOMP remains accessible to the BAM complex. In energetic terms, the enthalpy–entropy compensations consent with the binding modes of the two chaperones to maintain their interaction with uOmpX energetically favorable. Lastly, both chaperones synergistically disaggregate OmpX aggregates, suggesting additional interactions between both chaperones that enhance the disaggregation. We hypothesize that the tightly regulated multi-faceted functionalities of both chaperones enable cellular vitality to be maintained and regulated under normal and stress conditions.

## Methods

### Protein production

A tag-free double-cysteine variant of OmpX (OmpX_1,149_) without signal sequence was produced as inclusion bodies in *Escherichia coli* BL21(DE3) cells and purified following standard procedures including anion exchange chromatography, as previously reported^70,71^. After refolding, the protein was labeled with FRET donor (ATTO532, Atto-Tec) and acceptor (Abberior STAR 635P, Abberior) dyes. Details are given in the Supplementary Information.

Skp and SurA were produced as N-terminal hexahistidine (His_6_) fusion proteins in *E. coli* BL21 (DE3) cells and purified using immobilized metal affinity chromatography under denaturing conditions, as previously reported^32,60^. The proteins were refolded prior to experiments. Details on the production and purification are given in the Supplementary Information.

### Sample preparation for single-molecule measurements

A multi-step dilution protocol, as shown in Fig. 1a, was followed to prepare samples for smFRET and FCS measurements. In both FCS and smFRET experiments, the labeled protein was present at a concentration of 20 pM in buffer containing 20 mM Tris-HCl (pH 8.0) and 150 mM NaCl. The remaining concentration of GdmCl and LDAO was 6 mM and 87.5 nM, respectively. For FCS experiments involving OmpX aggregates, as shown in Fig. 5a, the concentration of GdmCl and LDAO was 12 mM and 175 nM, after addition of 1 μM uOmpX in the same buffer as used for smFRET experiments. Details are given in the Supplementary Information.

### smFRET and FCS measurements

Experiments were carried out using a single-molecule confocal fluorescence microscope as previously described^72^ and detailed in the Supplementary Information. Data analysis was performed with custom-written Matlab scripts (Mathworks). Single-molecule events were identified from the acquired photon stream by a burst search algorithm. Details about smFRET and FCS analysis procedures, including burst selection, data reduction, calculation of FRET efficiencies and fluorescence, are given in the Supplementary Information.

## Acknowledgements

We thank Sandro Keller (University of Graz) and all members of the Schlierf group for fruitful discussions. This work was supported by the TU Dresden institutional funds (M.S.), German Federal Ministry of Education and Research BMBF with grant 03Z2EN11 (M.S.), the Deutsche Forschungsgemeinschaft with SCHL 1896/3-1 and SCHL 1896/4-1 (M.S.), and Dresden International Graduate School for Biomedicine and Bioengineering (DFG GS97) to N.C.. G.K. acknowledges support by the European Research Council (ERC) under the European Union’s Horizon 2020 Framework Programme through the Marie Skłodowska-Curie grant MicroSPARK (agreement no. 841466) (G.K.), the Herchel Smith Funds of the University of Cambridge, and the Wolfson College Junior Research Fellowship.

## Author Contributions

N.C., A.H., G.K., and M.S. designed research; N.C. and A.H. carried out experiments; M.Q.M., G.K., and N.C. provided reagents; N.C. and A.H. analyzed data; A.H. implemented and developed analysis methods; N.C. and G.K. wrote initial draft; N.C., A.H., G.K., and M.S. edited and contributed to the final draft. M.S. supervised the research and acquired funding.

