## Supplementary Information for "Chaperones Skp and SurA dynamically expand unfolded outer membrane protein X and synergistically disassemble oligomeric aggregates"

### Supplementary figures

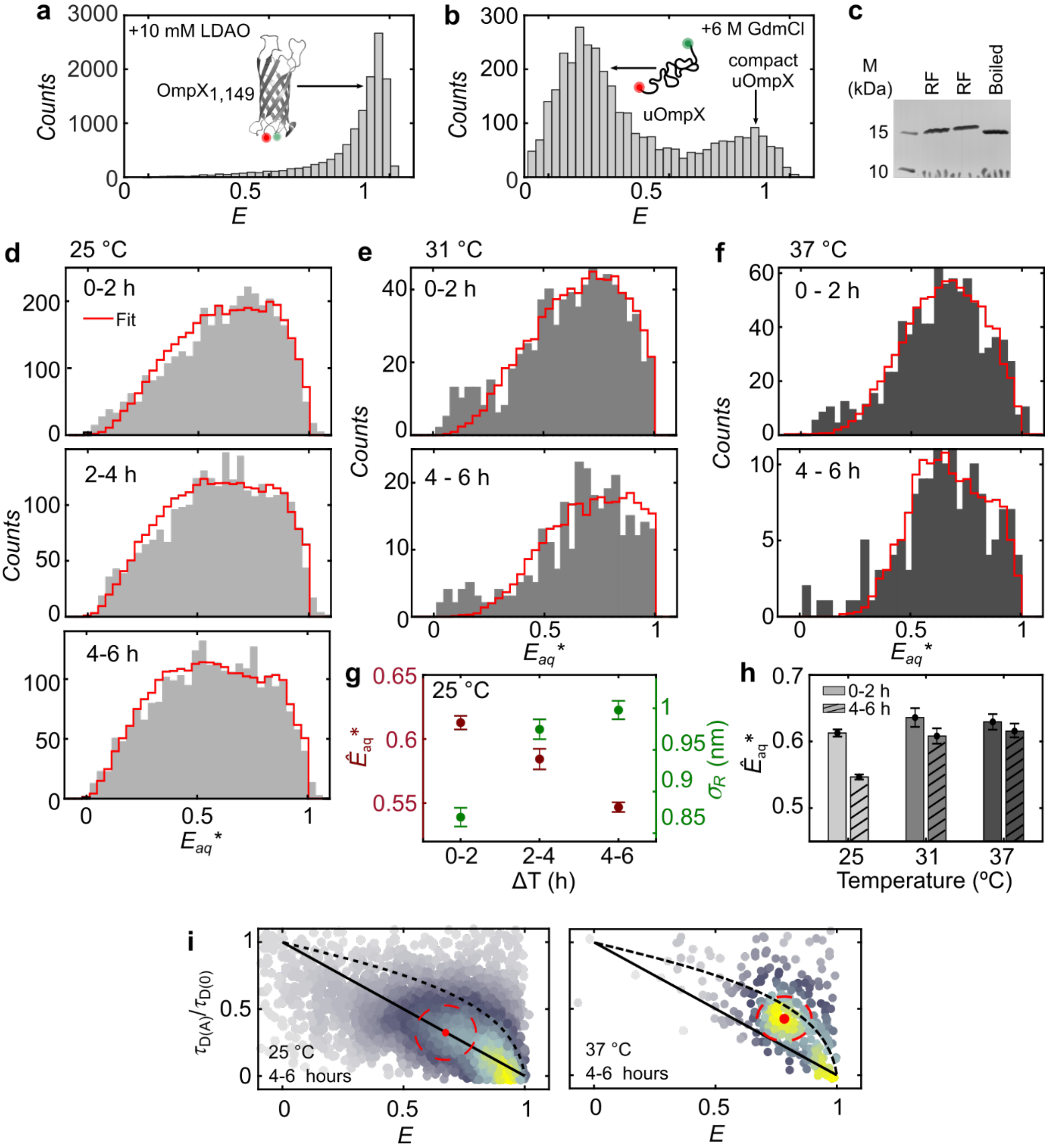


**Supplementary Figure 1. Refolded, unfolded, and aqueous OmpX (a)** FRET efficiency (*E*) histogram of refolded $OmpX_{1,149}$ in the presence of 10 mM LDAO. **(b)** *E* histogram of unfolded $OmpX_{1,149}$ in the presence of 6 M GdmCl without detergent. **(c)** SDS PAGE gel showing refolded OmpX fractions obtained after size exclusion chromatography. The boiled sample migrates differently than the folded protein and is shown for reference. The molecular weights of a marker (M) are indicated. **(d–f)** Fits (shown as red cityscapes) obtained from a two-state probability distribution analysis (PDA) taking into account the $uOmpX_{aq}$ and $uOmpX_{compact}$ state for **(d)** the first two hours (0–2 h), the next two hours (2–4 h), and the last two hours (4–6 h) at 25 °C, **(e)** the first two hours (0–2 h) and the last two hours (4–6 hours) at 31 °C, and **(f)** the first two hours (0–2 h) and the last two hours (4–6 h) at 37 °C. The analysis was done as described previously.[1] It allowed extraction of the mean apparent FRET efficiency,

$\hat{E}_{aq}$*, and the underlying width of the distance distribution, $\sigma_R$, of the $uOmpX_{aq}$ and $uOmpX_{compact}$ state of OmpX from the apparent FRET efficiency histograms for each time interval as given in Supplementary Table 2. The red line shows the fit for each PDA fit. **(g)** Average apparent FRET efficiency ($\hat{E}_{aq}$*) (maroon points) and the width of the underlying distance distribution ($\sigma_R$) (green points) with error bars, plotted against time. **(h)** $\hat{E}_{aq}$* of $uOmpX_{aq}$ state corresponding to 0–2 h and 4–6 h is plotted against the measurement temperature. **(i)** 2D scatter plot of the relative fluorescence lifetime of donor ($\tau_{D(A)}/\tau_{D(0)}$) vs. FRET efficiency ($E$) for the $OmpX_{1,149}$ measurement at 25 °C and 37 °C for 4–6 h. The solid line depicts the static FRET line and the dashed line indicates the expected correlation for a Gaussian chain. The red dot and the red dashed circle denote the center position and 68% area of the $uOmpX_{aq}$ population.

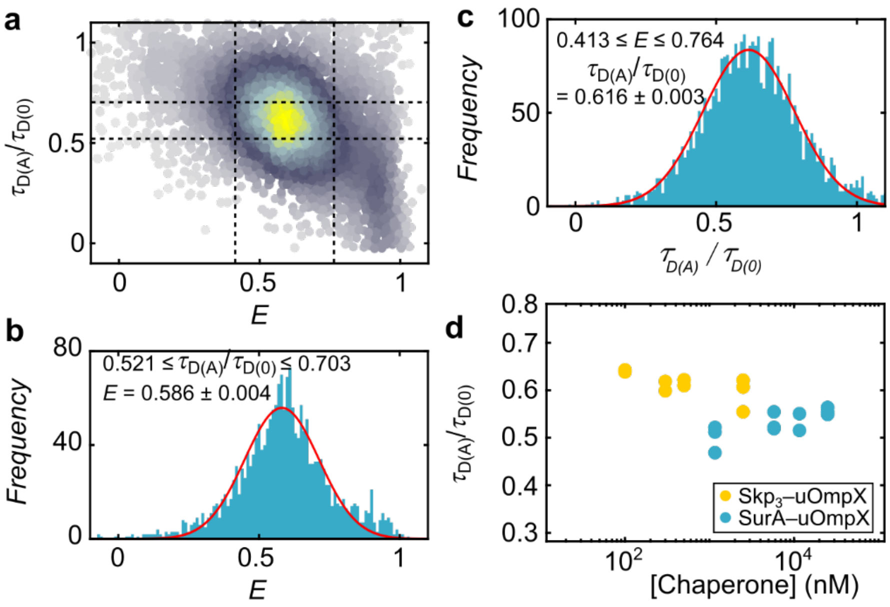


**Supplementary Figure 2. Workflow to obtain peak positions of *E* and ($\tau_{D(A)}/\tau_{D(0)}$). (a)** 2D scatter plot of the relative lifetime of donor ($\tau_{D(A)}/\tau_{D(0)}$) vs FRET Efficiency (*E*) with black dashed lines indicating the range of filters applied to obtain the center positions corresponding to **(b)** the FRET efficiency (*E*) and **(c)** $\tau_{D(A)}/\tau_{D(0)}$ as reported in the respective figures. **(d)** Center positions of $Skp_3$- (yellow spheres) and SurA-bound (blue spheres) uOmpX at all three temperatures (25 ˚C, 31 ˚C and 37 ˚C)). Note that the center positions for chaperone-bound species were calculated only for measurements which had a significant fraction of the chaperone–uOmpX complex (i.e., >100 nM [Skp3] at 25 ˚C and 31 ˚C and >2.5 μM [$Skp_3$] at 37 ˚C and >1.16 μM [SurA] at all three temperatures except for 11.6 μM [SurA] at 25 ˚C).

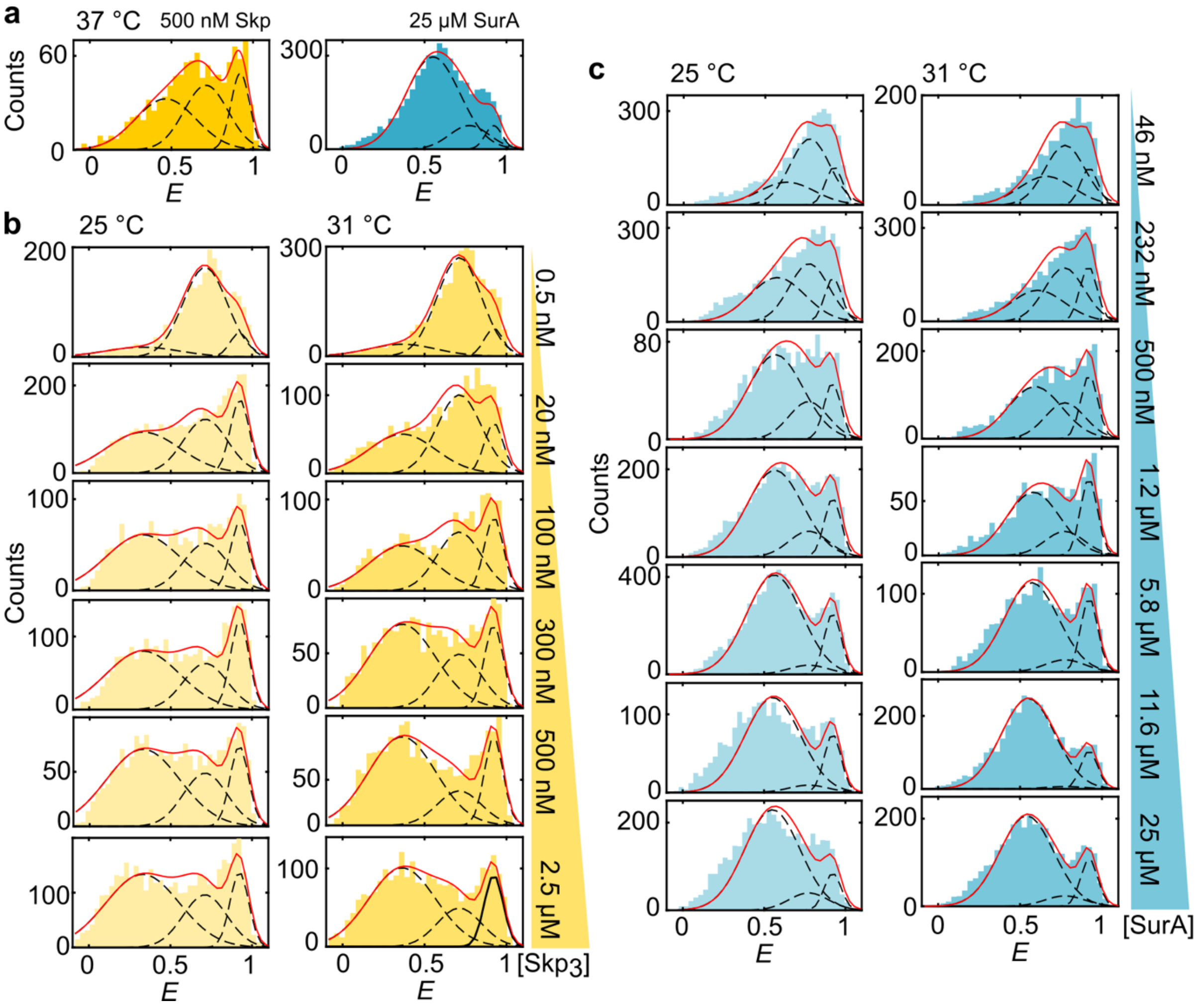


**Supplementary Figure 3. FRET efficiency histograms of chaperone-complexed OmpX. (a)** uOmpX in the presence of 500 nM $Skp_3$ and 25 µM SurA at 37 °C, **(b)** uOmpX complexed with increasing $Skp_3$ concentration at 25 °C and 31 °C, **(c)** uOmpX complexed with increasing SurA concentration at 25 °C and 31 °C.

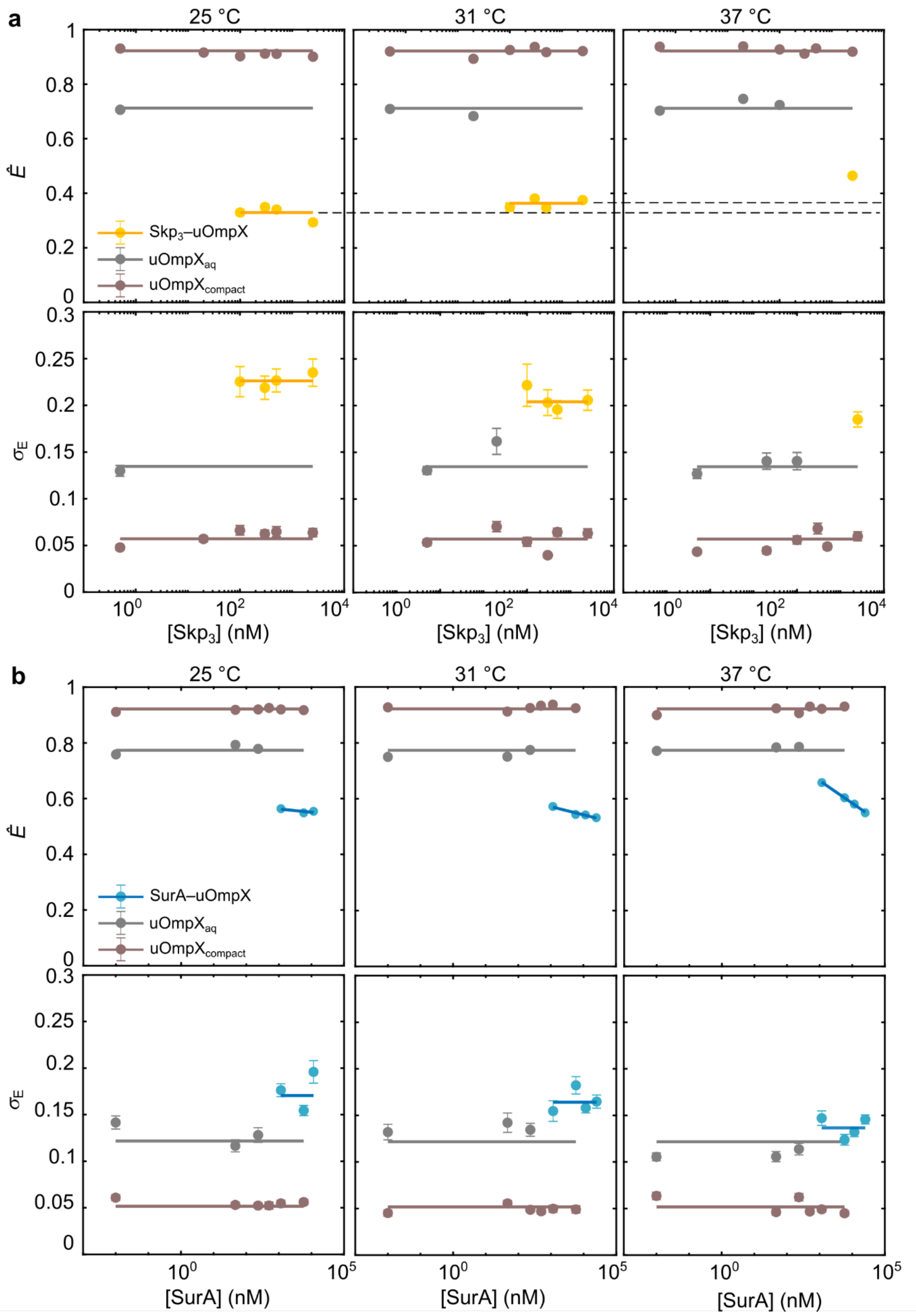


**Supplementary Figure 4. Trend in peak position and width of FRET efficiency distributions in presence of chaperones of the uOmpX$_{aq}$, uOmpX$_{compact}$, and chaperone-bound OmpX.** **(a)** Average FRET efficiencies of the three states are plotted against Skp$_3$ concentration. While the compact (brown) and unbound FRET (grey) states and their widths ($\sigma_E$) do not change with temperature or concentration, the peak position ($\hat{E}$) of the bound state (yellow) decreases with increase in temperature, **(b)** Peak positions, $\hat{E}$, are plotted against SurA concentration. While the uOmpX$_{compact}$ (brown) and the uOmpX$_{aq}$ FRET (grey) states and their distance widths ($\sigma_E$) do not change with temperature or concentration, the peak position ($\hat{E}$) of the SurA–uOmpX state (blue) increases with concentration at each temperature. The fit $\hat{E}$ and $\sigma_E$ are reported in Supplementary Table 3.

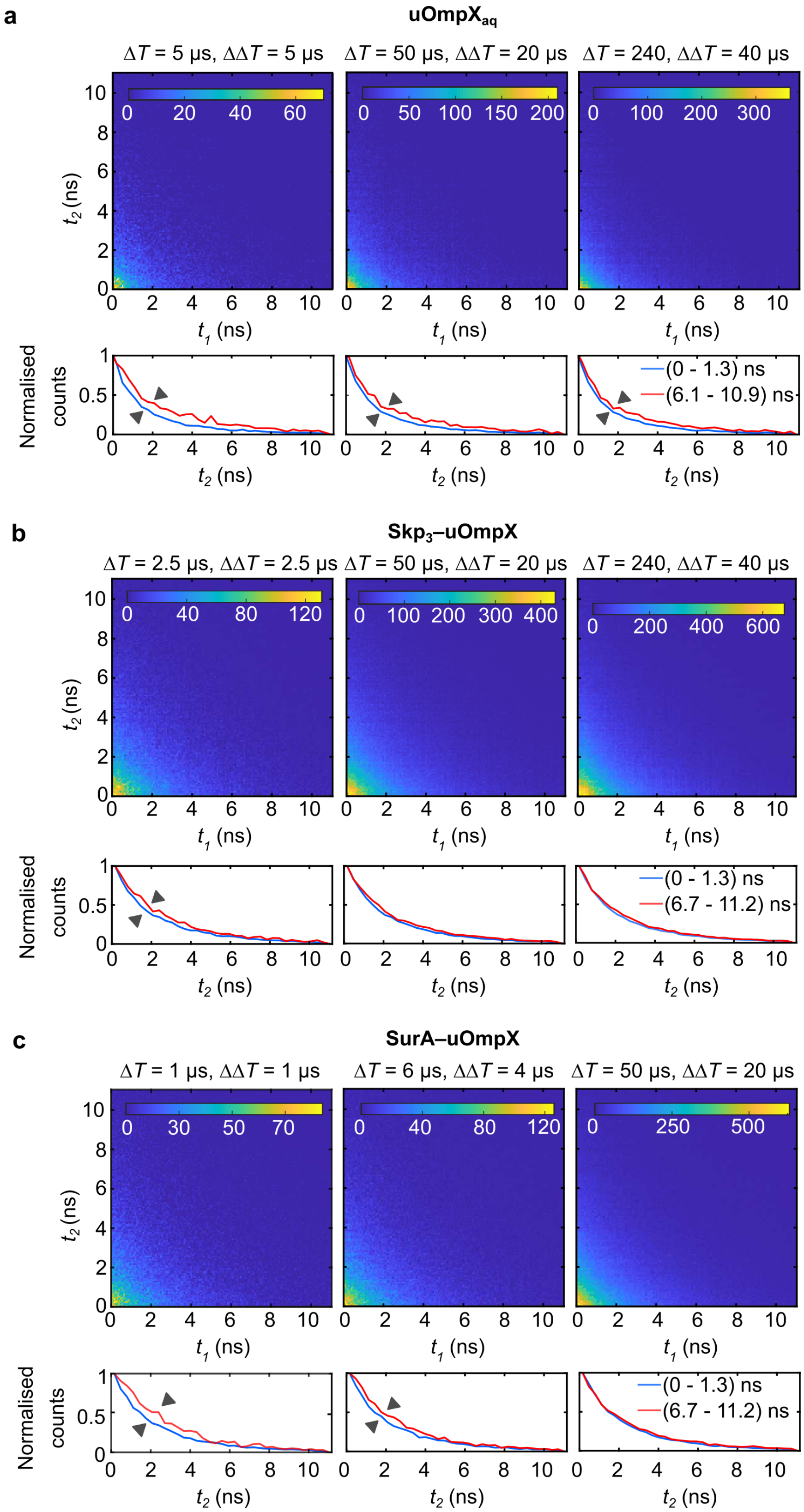


**Supplementary Figure 5. Species-filtered 2D emission delay correlation maps and 1D emission-delay histograms of increasing time intervals (Δ*T*) with ΔΔ*T* as the window size. (a)** For $uOmpX_{aq}$ fraction. **(b)** For the $Skp_3$-bound uOmpX fraction at [$Skp_3$] = 2.5 μM **(c)** For the SurA-bound uOmpX fraction at [SurA] = 11.6 μM.

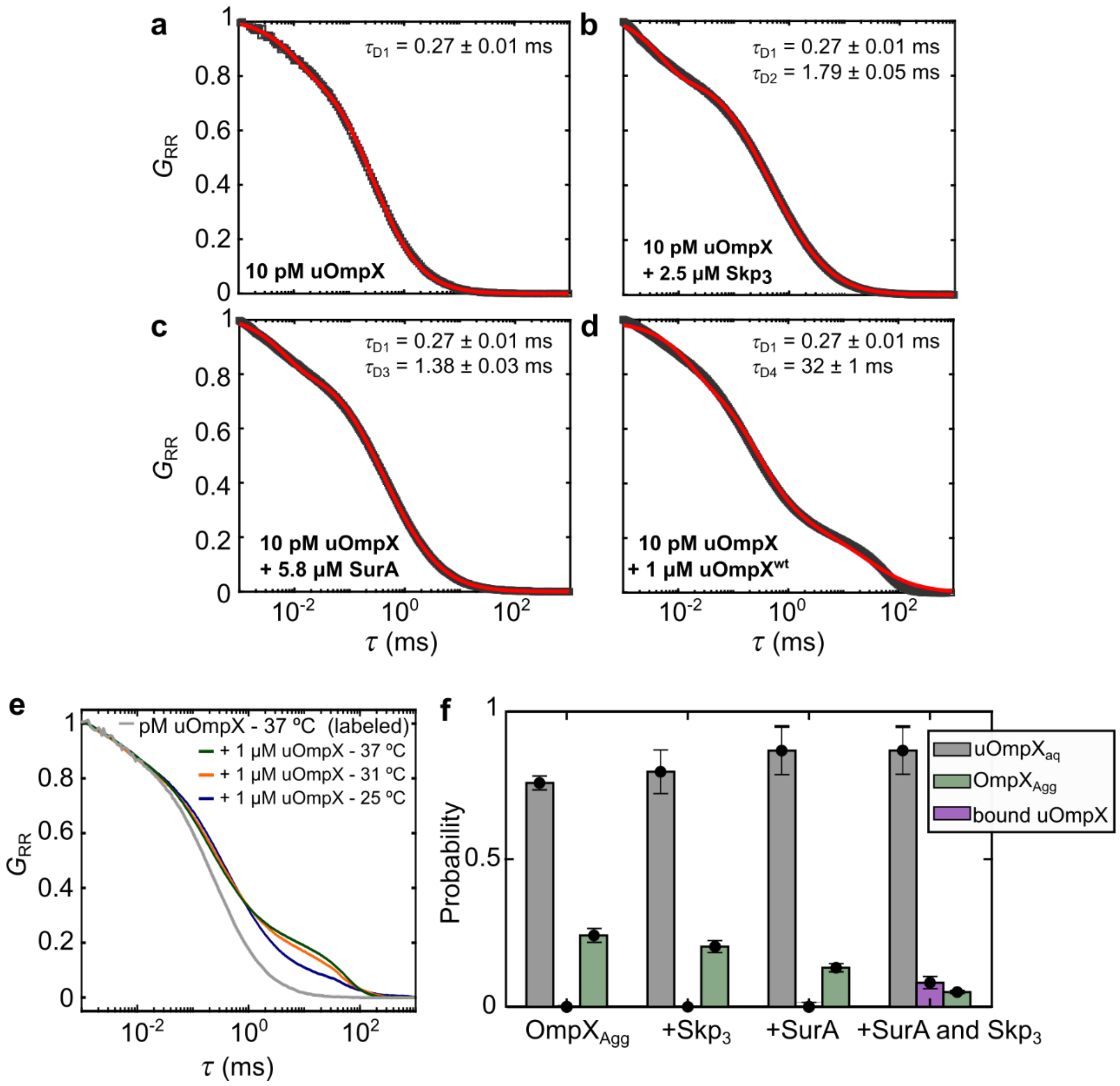


**Supplementary Figure 6. FCS analysis of OmpX$_{Agg}$ disaggregation by Skp$_3$ and SurA**. **(a)** Aqueous OmpX (uOmpX$_{aq}$). **(b)** Skp$_3$ + uOmpX. **(c)** SurA + uOmpX. **(d)** 10 pM labeled uOmpX + 1 µM unlabeled uOmpX in aqueous buffer. $\tau_{Di}$ is the individual diffusion time of each state *i* and $G_{RR}$ is the autocorrelation function for the acceptor photons. The data are shown as black squares and the fits as a red line. **(e)** FCS curves of 10 pM labeled uOmpX at 37 °C and 10pM labeled uOmpX + 1µM unlabeled uOmpX in aqueous buffer at 37 °C (green), 31 °C (orange) and 25 °C (dark blue). The aggregation decreases with decreasing temperature, yet a significant fraction of OmpX$_{Agg}$ is observed at each temperature. **(f)** Fraction of uOmpX$_{aq}$ (grey bar), OmpX$_{Agg}$ (green), and chaperone-bound uOmpX (purple).

### Supplementary Tables

**Supplementary Table 1. Fluorescence lifetimes and time-resolved anisotropies of $uOmpX_{aq}$ and $Skp_3$- and SurA-bound uOmpX.** The fluorescence lifetime in the absence of the acceptor ($\tau_{D(0)}$) is, within error, constant among the different measurements, implying a constant quantum yield of the donor fluorophore. Because the rotational correlation times of the donor and acceptor ($\rho_{GG,fast}$) and ($\rho_{RR,fast}$), respectively, are smaller than the minimal time of energy transfer ($1/k_{FRET}$), and because the combined anisotropy ($r_C$) is smaller than 0.2, a sufficient rotational averaging of the dipoles ($\kappa^2$ = 2/3) can be assumed.

| | $\tau_{D(0)}$ (ns) | $\tau_{D(A)}$ (ns) | ($1/k_{FRET}$) (ns) | $\rho_{GG,fast}$ (ns) | $\rho_{RR,fast}$ (ns) |
|---|---|---|---|---|---|
| **$uOmpX_{aq}$** | 3.278 ± 0.004 | 1.18 ± 0.02 | 1.85 ± 0.04 | 0.54 ± 0.27 | 1.54 ± 0.79 |
| **$Skp_3$–uOmpX** | 3.195 ± 0.003 | 2.08 ± 0.01 | 5.98 ± 0.07 | 0.80 ± 0.29 | 1.87 ± 0.77 |
| **SurA–uOmpX** | 3.212 ± 0.002 | 1.96 ± 0.01 | 5.04 ± 0.04 | 1.23 ± 0.32 | 1.19 ± 0.36 |

| | $r_{\infty,GG}$ | $r_{\infty,GR}$ | $r_{\infty,RR}$ | $r_C$ |
|---|---|---|---|---|
| **$uOmpX_{aq}$** | 0.08 ± 0.02 | 0.018 ± 0.004 | 0.13 ± 0.04 | 0.10 ± 0.02 |
| **$Skp_3$–uOmpX** | 0.13 ± 0.02 | 0.032 ± 0.004 | 0.14 ± 0.03 | 0.14 ± 0.02 |
| **SurA–uOmpX** | 0.08 ± 0.02 | 0.019 ± 0.003 | 0.17 ± 0.02 | 0.11 ± 0.02 |

**Supplementary Table 2**. The probability of the uOmpX$_{aq}$ state ($p_{aq}$), apparent FRET efficiencies of uOmpX$_{aq}$ ($E_{aq}$*) and uOmpX$_{compact}$ state ($E_{compact}$*) and the respective distance widths $\sigma_{aq}$ and $\sigma_{compact}$ obtained from the two state PDA used to show expansion of the bound state on a timescale of hours at 25 °C. The best reduced chi-square ($\chi^2$) is given.

| **Δ*t* (h)** | $p_{aq}$ | $E_{aq}$* | $E_{compact}$* | $\sigma_{aq}$ **(nm)** | $\sigma_{compact}$ **(nm)** | $\chi^2$ |
|---|---|---|---|---|---|---|
| **0–2** | 0.953 ± 0.014 | 0.613 ± 0.005 | 0.857 ± 0.002 | 0.869 ± 0.011 | 0.104 ± 0.236 | 5.39 |
| **2–4** | 0.974 ± 0.013 | 0.584 ± 0.008 | 0.857 ± 0.002 | 0.975 ± 0.012 | 0.104 ± 0.236 | 3.06 |
| **4–6** | 0.966 ± 0.012 | 0.547 ± 0.003 | 0.857 ± 0.002 | 0.998 ± 0.011 | 0.104 ± 0.236 | 1.91 |

**Supplementary Table 3**. Peak position of each FRET state (chaperone bound: $\hat{E}_b$, unbound or aqueous: $\hat{E}_{aq}$ and compact: $\hat{E}_c$) and their widths (bound: $\sigma_{E\text{-}b}$, uOmpX$_{aq}$: $\sigma_{E\text{-}aq}$ and uOmpX$_{compact}$: $\sigma_{E\text{-}c}$) for both Skp and SurA interaction.

| Chaperone | *T* | $\hat{E}_b$ | $\hat{E}_{aq}$ | $\hat{E}_c$ | $\sigma_{E\text{-}b}$ | $\sigma_{E\text{-}aq}$ | $\sigma_{E\text{-}c}$ |
|---|---|---|---|---|---|---|---|
| Skp | 25 °C | 0.33 ± 0.02 | 0.71 ± 0.01 | 0.922 ± 0.003 | 0.226 ± 0.005 | 0.135 ± 0.008 | 0.057 ± 0.002 |
| | 31 °C | 0.36 ± 0.02 | 0.71 ± 0.01 | 0.922 ± 0.003 | 0.204 ± 0.008 | 0.135 ± 0.008 | 0.057 ± 0.002 |
| | 37 °C | 0.47 ± 0.01 | 0.71 ± 0.01 | 0.922 ± 0.003 | 0.185 ± 0.008 | 0.135 ± 0.008 | 0.057 ± 0.002 |
| SurA | 25 °C | -0.005×log([SurA])+0.598 | 0.774 ± 0.006 | 0.922 ± 0.002 | 0.17 ± 0.02 | 0.122 ± 0.006 | 0.052 ± 0.001 |
| | 31 °C | -0.013×log([SurA])+0.659 | 0.774 ± 0.006 | 0.922 ± 0.002 | 0.164 ± 0.009 | 0.122 ± 0.006 | 0.052 ± 0.001 |
| | 37 °C | -0.350×log([SurA])+0.906 | 0.774 ± 0.006 | 0.922 ± 0.002 | 0.1366 ± 0.009 | 0.122 ± 0.006 | 0.052 ± 0.001 |

**Supplementary Table 4.** Thermodynamic parameters: Enthalpic change ($\Delta H$) and entropic change ($\Delta S$) are obtained for $Skp_3$ and SurA interaction with uOmpX through global $\chi^2$ minimization. $c_{1/2\text{-}25\ °C}$, $c_{1/2\text{-}31\ °C}$ and $c_{1/2\text{-}37\ °C}$ denotes the half association concentration at 25 °C, 31 °C and 37 °C respectively with $n$ as the hill coefficient. $f_c$(25 °C), $f_c$(31 °C) and $f_c$(37 °C) are the fractions of $uOmpX_{compact}$ state at 25 °C, 31 °C and 37 °C and $f_{b\text{-}max}$(25 °C), $f_{b\text{-}max}$(31 °C), $f_{b\text{-}max}$(37 °C) is the maximum fraction of chaperone-bound uOmpX at 25 °C, 31 °C and 37 °C respectively. It was the same for SurA at all temperatures. The free energy of interaction so obtained at each temperature are also reported: $\Delta G_{25\ °C}$, $\Delta G_{31\ °C}$ and $\Delta G_{37\ °C}$.

| **Chaperone** | ***n*** | **$\Delta H$ (kJ mol$^{-1}$)** | **$\Delta S$ (kJ mol$^{-1}$ K$^{-1}$)** | **$f_c$(25 °C)** | **$f_c$(31 °C)** | **$f_c$(37 °C)** | **$f_{b,max}$(25 °C)** |
|---|---|---|---|---|---|---|---|
| SurA | **1.0** | –75 ± 12 | –0.12 ± 0.04 | 0.118 ± 0.009 | 0.14 ± 0.01 | 0.114 ± 0.008 | 0.92 ± 0.01 |
| SurA | **1.5** | –132 ± 24 | –0.24 ± 0.08 | 0.119 ± 0.008 | 0.141 ± 0.009 | 0.111 ± 0.008 | 0.88 ± 0.01 |
| $Skp_3$ ([Skp]/3) | **1.0** | –298 ± 19 | –0.84 ± 0.06 | 0.195 ± 0.003 | 0.184 ± 0.007 | 0.179 ± 0.006 | 0.69 ± 0.01 |
| $Skp_3$ [2] | **1.0** | –480 ± 21 | –1.42 ± 0.07 | 0.195 ± 0.004 | 0.188 ± 0.005 | 0.183 ± 0.006 | 0.70 ± 0.01 |

| **$f_{b,max}$(31 °C)** | **$f_{b,max}$(37 °C)** | **$c_{1/2\text{-}25\ °C}$ (nM)** | **$c_{1/2\text{-}31\ °C}$ (nM)** | **$c_{1/2\text{-}37\ °C}$ (nM)** | **$\Delta G_{25\ °C}$ (kJ mol$^{-1}$)** | **$\Delta G_{31\ °C}$ (kJ mol$^{-1}$)** | **$\Delta G_{37\ °C}$ (kJ mol$^{-1}$)** | $\chi^2$ |
|---|---|---|---|---|---|---|---|---|
| 0.92 ± 0.01 | 0.92 ± 0.01 | 109.67 ± 0.01 | 199.06 ± 0.02 | 356.05 ± 0.06 | –39.7 ± 0.3 | –39.0 ± 0.3 | –38.3 ± 0.4 | 3.83 |
| 0.88 ± 0.01 | 0.88 ± 0.01 | 102.89 ± 0.02 | 205.94 ± 0.02 | 407.40 ±0.07 | –59.9 ± 0.6 | –58.4 ± 0.4 | –56.9 ± 0.6 | 3.97 |
| 0.81 ± 0.02 | 0.998 ± 0.004 | 3.554 ± 0.001 | 36.993 ± 0.005 | 358.88 ± 0.02 | –48.3 ± 0.7 | –43.3 ± 0.3 | –38.3 ± 0.2 | 3.19 |
| 0.77 ± 0.01 | 0.98 ± 0.05 | 0.088 ± 0.001 | 3.942 ± 0.001 | 154.39 ± 0.03 | –57.5 ± 0.9 | –49.0 ± 0.7 | –40.5 ± 0.7 | 3.46 |

**Supplementary Table 5.** FCS results of aqueous OmpX ($OmpX_{aq}$) i.e., without chaperones, SurA–uOmpX, or $Skp_3$–uOmpX and aggregated OmpX ($OmpX_{Agg}$). *Q* is the brightness, $\tau_D$ the diffusion time, $\tau_T$ is the characteristic triplet time and *T* is the triplet fraction.

| **Measurement** | ***Q* (kHz)** | **$\tau_D$ (ms)** | **$\tau_T$ (µs)** | ***T*** |
|---|---|---|---|---|
| $OmpX_{aq}$ | 48.79 | 0.27 ± 0.01 | 9.3 ± 0.4 | 0.164 ± 0.003 |
| $Skp_3$–uOmpX | 47.85 (-0.1<E<0.6) | 1.79 ± 0.05 | 4.4 ± 0.1 | 0.223 ± 0.001 |
| SurA–uOmpX | 52.82 (-0.1<E<0.75) | 1.38 ± 0.03 | 5.9 ± 0.1 | 0.191 ± 0.001 |
| $OmpX_{Agg}$ | 52.66 | 32 ± 1 | 1.5 ± 0.1 | 0.185 ± 0.001 |

### Supplementary Methods

**Preparation of fluorescently labeled OmpX.** The mature sequence of *Escherichia coli* OmpX (residues 24–171) was cloned from genomic DNA into a pET28a expression vector between NcoI and XhoI restriction sites, yielding a construct encoding OmpX without its signal peptide and without any affinity tags. The QuikChange Lightning Multi Site-Directed Mutagenesis Kit (Agilent) was used to introduce two cysteine residues, one cysteine at the first position (A1C) of the OmpX sequence and the second cysteine C-terminally of the OmpX chain (149C). This resulted in a plasmid encoding a 149-amino-acid-residue double-Cys variant denoted as $OmpX_{1,149}$.

$OmpX_{1,149}$ was produced as inclusion bodies (IBs) following previously established protocols.[3–5] Briefly, the plasmid encoding $OmpX_{1,149}$ was transformed into *E. coli* BL21 (DE3) cells (Agilent). Cells were grown at 37 °C in Luria-Bertani (LB) medium containing 50 µg $ml^{-1}$ kanamycin (Carl Roth) to $OD_{600}$ = 0.6. Expression was induced by 0.4 mM isopropyl-β-d-thiogalactopyranoside (IPTG; Roth). Cells were harvested after overnight induction at 20 °C, resuspended in lysis buffer (50 mM Tris-HCl (pH 8.0), 40 mM EDTA, 25% (*w*/*v*) sucrose) and lysed using an EmulsiFlex-C3 (Avestin) high-pressure homogenizer. The lysate was centrifuged for 45 min at 7'000 × *g* and 4 °C to pellet IBs and remove the soluble supernatant. Purification from IBs was done as described previously[3–5]. Briefly, IBs were solubilized with 8 M urea (Sigma-Aldrich) overnight on a rotator at 4 °C and unfolded $OmpX_{1,149}$ in the solubilized fraction was separated from residual impurities by centrifugation for 1 h at 7'000 × *g* and 4 °C. Refolding was carried out at 50 °C by drop dilution of unfolded $OmpX_{1,149}$ in the refolding buffer (20 mM Tris-HCl, 2 mM EDTA (pH 8.3) containing 35 mM LDAO) for 16 h under stirring. The folded protein was further isolated with anion exchange chromatography using a HiTrap DEAE FF 1 ml column (GE Healthcare).

The purified protein was site-specifically labeled via thiol-maleimide chemistry with 20-fold excess of maleimide-functionalized donor (ATTO532; Atto-Tec) and acceptor (Abberior STAR 635P; Abberior) fluorophores in buffer containing 20 mM Tris-HCl (pH 7.2), 2 mM EDTA, and 17.5 mM LDAO. The labeled protein was separated from unbound dyes by size-exclusion chromatography using a Superdex 75 10/300 GL column (GE Healthcare). The purified protein was then stored at 4 °C for immediate use.

**Preparation of Skp and SurA.** The mature sequences of *E. coli* Skp and SurA (Skp: residues 21–161); SurA: residues 21–428) were cloned from genomic DNA into pET28a expression vectors between NdeI and XhoI restriction sites, yielding constructs encoding Skp and SurA without their respective signal peptides and an N-terminal hexahistidine ($His_6$) tag.

Expression and purification were performed as previously described[6,7]. Briefly, the plasmids encoding $His_6$-Skp and $His_6$-SurA were transformed into *E. coli* BL21 (DE3) cells. Cells were grown at 37 °C in LB medium containing 50 µg ml$^{-1}$ kanamycin to $OD_{600}$ = 0.6. Expression was induced by 0.4 mM IPTG. Cells were harvested after overnight induction, resuspended in lysis buffer (150 mM Tris-HCl (pH 7.2), 20 mM NaCl, 20 mM imidazole; supplemented with an EDTA-free protease inhibitor cocktail (cOmplete, Roche)) and lysed using an EmulsiFlex-C3 high-pressure homogenizer. The cell lysate was cleared by centrifugation at 18'000 × *g* for 30 min at 4 °C. The supernatant, containing Skp or SurA, was denatured with 6 M GdmCl (Thermo Scientific) overnight on a rotator and then applied to nickel-nitrolotriacetic acid (Ni-NTA) columns (HisTrap HP 5 ml; GE Healthcare). Elution of protein was done with 150 mM Tris-HCl (pH 7.2), 20 mM NaCl, 500 mM imidazole, and 6 M GdmCl in a gradient. Both proteins eluted at ~50% imidazole gradient. The pooled fractions were dialyzed overnight at 4 °C in assembly buffer (20 mM Tris-HCl (pH 7.2), 150 mM NaCl) and then stored at 4 °C for immediate use or at –80 °C for long term storage.

**Preparation of chaperone–uOmpX complexes for single-molecule measurement.** A multi-step dilution protocol, as shown in Fig. 1a was followed to prepare samples for smFRET measurements. Accordingly, refolded and labelled $OmpX_{1,149}$ was first denatured at 1 µM for at least 24 h with unfolding buffer (20 mM Tris-HCl (pH 8.0), 6 M GdmCl, 2 mM EDTA) resulting in unfolded $OmpX_{1,149}$ (uOmpX). The next dilution to 20 nM was performed again in unfolding buffer. uOmpX was diluted to its final measurement concentration of 20 pM in the measurement buffer (20 mM TrisHCl and 150 mM NaCl, pH 8.0). This final dilution was done in the absence or presence of the chaperone ($Skp_3$ or SurA) at a concentration as mentioned in the results section. At this stage, the concentration of LDAO and GdmCl was 6 mM and 0.06 mM respectively.

**smFRET measurements.** Experiments were carried out using a custom-built setup combining confocal single-molecule spectroscopy with time-correlated single-photon counting (TCSPC), pulsed interleaved excitation (PIE) and fluorescence anisotropy detection for multiparameter-fluorescence detection (MFD)[8]. Donor and acceptor fluorophores were excited with linearly polarized 530-nm and 640-nm picosecond pulsed lasers (LDH-P-FA-530L and LDH-D-C-640, PicoQuant) driven in PIE mode at a total repetition rate of 50 MHz. The laser beams were coupled into a polarization-maintaining single-mode optical fiber (P3-488PM-FC-2, Thorlabs), collimated (60FC-T-4-RGBV42-47, Schäfter+Kirchhoff), and focused by a water immersion objective (CFI Plan Apo WI 60x, NA 1.2, Nikon). Emitted fluorescent light was collected by the same objective, separated from the excitation light by a dual-edge dichroic mirror (zt532/642rpc, Chroma), and focused on a 50-µm pinhole (Thorlabs). Donor and acceptor photons were spectrally separated by single-edge dichroic mirrors (FF650-Di01, Semrock) after a polarizing beam splitter (CM1-PBS251, Thorlabs), band-pass-filtered (FF01-582/75, Semrock; ET700/75M, Chroma), and focused onto four single-photon-counting avalanche diodes (SPCM-AQR, Excelitas). Photons were registered by four individual TCSPC modules (Hydra Harp, PicoQuant) with a time resolution of 16 ps. Synchronization with the lasers for alternating excitation was accomplished with the aid of a diode laser driver (PDL828, PicoQuant).

Measurements were performed in custom-built sample chambers on freely diffusing molecules by placing the confocal volume into solution at an axial position ca. 60 µm above the surface of the cover slide. The sample chamber was passivated with bovine serum albumin (Sigma). The illumination power was 110 µW for excitation of the donor dye and 90 µW for direct excitation of the acceptor dye measured before the objective. During all measurements, the temperature of the sample chamber was controlled by an objective collar connected to a refrigerated/heated circulator (F25-MC, Julabo, Germany). Measurements were performed at ~10 pM concentration of labelled protein to maintain single molecule conditions at appropriate burst rates.

Data analysis was performed with custom-written Matlab scripts (Mathworks). Briefly, single-molecule events were identified from the acquired photon stream using a burst search algorithm. Only bursts with a maximum inter-photon time of 50 µs and a minimum total number of 40 photons were selected. The FRET efficiency ($E$) of each single-molecule burst was calculated according to $E = n_A/(n_D+n_A)$, where $n_D$ and $n_A$ are the number of photons detected in the donor and acceptor channel after donor excitation, respectively, and corrected for background, acceptor direct excitation ($\alpha$ = 0.0881), spectral cross-talk ($\beta$ = 0.0247) and differences in detection efficiencies, and quantum yields ($\gamma_{25\,°C}$ = 0.4567, $\gamma_{31\,°C}$ = 0.4481, $\gamma_{37\,°C}$ = 0.4390) as described previously[8]. Only molecules exhibiting a stoichiometry ratio of $S$ between 0.2 and 0.75 and an alternating laser excitation−two-channel kernel-based density distribution estimator (ALEX-2CDE) score smaller than 12 were selected for further analysis. Here, $S = n^{Dexc}/(n^{Dexc}+n^{Aexc})$, where $n^{Dexc}$ is the total number of corrected counts after donor excitation and $n^{Aexc}$ is the total number of corrected counts after acceptor excitation. Additionally, to minimize the number of bursts resulting from fluorescence background (e.g., impurities) and multiple molecule events that display signals in the donor and acceptor channels, bursts were rejected if the ratio of the number of donor photons to number of acceptor photons after donor excitation was more than 0.75 and 0.9 for measurements with SurA and Skp, respectively. The remaining bursts were analyzed to construct FRET efficiency histograms with bin widths of 0.033. The donor lifetime of each burst was estimated by the average microtime of donor photons subtracted by the average delay time of the IRF. The corresponding lifetime of the donor in absence of any acceptor was extracted from the donor only population with $S$ > 0.95.

**Analysis of population heterogeneity.** First, the center positions of $Skp_3$– and SurA–uOmpX distributions were extracted as illustrated in Supplementary Fig. 2 by Gaussian fitting. In a next step, the resulting coordinates were used to analyze the underlying inter-dye distance distribution showing interconversion dynamics on the microsecond timescale. To this end, the inter-dye distance distribution was modelled with a log-normal probability density function:

$$\mathrm{p}(R)=\frac{1}{\sqrt{2\pi}\sigma_\mathrm{R}R}\exp\left(-\frac{\left(\ln(R)-\mu_R\right)^2}{2\sigma_R{}^2}\right) \qquad (1)$$

with $\sqrt{e^{\sigma_R^2}-1}$ being the coefficient of variance, CV, and $e^{\mu_R+\frac{1}{2}\sigma_R^2}$ being the expected distance of the distribution. A corresponding coordinate of the relative donor lifetime,

$\tau_{D(A)}/\tau_{D(0)}$, and FRET efficiency, $E$, is then obtained by integration over time and distance, respectively, as described by Soranno et. al[9]. In the global fit, for each variation of $\sigma_R$, a dynamic $\tau_{D(A)}/\tau_{D(0)}$ vs $E$ curve was calculated for the range $\mu_R$ = {0.1–20} nm so as to record the chi-squared value. Finally, the most likely width, $\sigma_R$, of the log-normal distribution corresponding to the minimal chi-squared were obtained for the unbound, Skp- and SurA-bound states of OmpX and corresponding CV values have been calculated.

**Species filtered Two-Dimensional Fluorescence Lifetime Correlation Spectroscopy (2D FLCS).** In a first step, only fluorescence bursts of $uOmpX_{aq}$, $Skp_3$–uOmpX or SurA–uOmpX were selected by FRET efficiency filtering (–0.1 < $E$ < 1.1; 0.15 < $E$ < 0.7 and 0.3 < $E$ < 0.8, respectively). Then, photon pairs of these FRET efficiency-filtered fluorescence bursts with a time gap matching the time interval of $\Delta T$ and a window of $\Delta\Delta T$ were sorted according to their initial ($t_1$) and final ($t_2$) microtime (photon delay time with respect to the excitation pulse) in the two-dimensional emission-delay histogram (Supplementary Fig. 5) as described[10,11]. In order to disentangle the timescale of interconversion dynamics between different molecular states slices of the 2D histogram with either short (< 1.3 ns) or long (> 6.1 ns) initial microtimes have been extracted. This lifetime filtering is equal to a molecular synchronization, where only similar conformations with comparable initial FRET efficiencies (Fig. 3a, second and third panel) are selected. The similarity of the two derived 1D emission-delay histograms of final microtimes ($t_2$) shows then the amount of equilibration in the selected time interval ($\Delta T$) and window ($\Delta\Delta T$). If the time interval $\Delta T$ is within the range of the timescale of the interconversion dynamics (between $\tau^1_{D(A)}$ and $\tau^2_{D(A)}$), a correlation between initial and final microtime becomes visible as a separation between the two 1D emission-delay histograms.

**Calculation of thermodynamic parameters.** First, the set of histograms derived at the three different temperatures and from increasing chaperone concentrations were globally fitted using the shared variables $\Delta H$, $\Delta S$, and $f_{b,max}(T)$ and the individual variable $f_c(T)$. During optimization for each pair of {$\Delta H$, $\Delta S$, $f_{b,max}(T)$ and $f_c(T)$} the corresponding association constants, $K_a(T)$, were calculated for the respective measurement temperature (Equation 1, main text) as illustrated in the van't Hoff plot in Fig. 4a. The resulting association constants were then used to determine the individual theoretical fraction of bound OmpX, $f_b$, for the respective measurement temperature and chaperone concentration using the Hill equation:

$$f_b = f_{b,max}(T)\frac{[Chaperone]^n}{[Chaperone]^n + K_d(T)} \qquad (2)$$

where $K_d=1/K_a$ denotes the dissociation constant, $f_{b,max}(T)$ the maximal bound fraction at temperature $T$, and $n$ the Hill coefficient. It should be noted that we have considered the trimeric Skp concentration simply by dividing the monomeric concentration by 3 as it was diluted from a high concentration stock (~75 μM) for complexing with uOmpX (when considering a slow dissociation kinetics, Skp trimers would not have dissociated by the time of dilution). A case where the trimeric concentration was chosen, according to the previously published data[2], is also shown in Supplementary Table 4. This assumption follows from the fact that the substrate uOmpX was added to the buffer containing chaperone before a trimer–monomer equilibrium was established. Subsequently, for every measurement condition a FRET distribution was calculated using the FRET parameters (center position and width of the individual FRET efficiency population) and the amplitudes $f_b(1-f_c(T))$, $(1-f_b)$ $(1-f_c(T))$ and $f_c(T)$ using the photon statistics of the measured fluorescence bursts (Monte Carlo Simulation). Finally, the overall reduced chi-squared was calculated from the residuum of the theoretical and measured FRET efficiency curves. By varying the optimization parameters $\{\Delta H, \Delta S, f_c(T), f_{b,max}(T)\}$ the reduced Chi-square was minimized to find the best estimators for the change in enthalpy and entropy as well as $f_c(T)$ and $f_{b,max}(T)$. The errors of the fitted thermodynamic parameters were derived from 200 boot strapping steps, where for each measurement condition FRET efficiency histograms were constructed from subset of randomly drawn bursts.

**Fluorescence correlation spectroscopy.** In order to quantify the fractions of OmpX aggregates in solution, first the diffusion time of uOmpX was characterized at low protein concentration (~10 pM) in the absence of $Skp_3$ and/or SurA, respectively. To this end, the autocorrelation function was calculated from the collected acceptor photons (red PIE pulse) and fitted by Eq. 3 (see Supplementary Fig. 6a–d).

$$G(t) = \frac{\sum_{i=1}^{k}\left(Q_i^2\right)F_i g_i(t)}{N\left(\sum_{i=1}^{k} Q_i F_i\right)^2}\left(1+\frac{T}{1-T}\exp\left(-\frac{\tau}{\tau_T}\right)\right) \qquad (3)$$

$$g_i(\tau) = \left(1+\frac{\tau}{\tau_{\mathrm{Diff}}}\right)^{-1}\left(1+\frac{\tau}{\kappa^2\tau_{\mathrm{Diff}}}\right)^{-\frac{1}{2}}$$

with $F_i$ and $\tau_{Diff}$ being the fraction and diffusion time of species *i*, respectively. The parameter *N* denotes the average number of molecules in the confocal volume. We assumed a similar triplet state fraction (*T*) and characteristic triplet time ($\tau_T$) in the bound and unbound state. *k* denotes the number of diffusing components. In the presence of chaperones two diffusion components (*k* = 2) were used to model the extracted correlation curve. The molecular brightness ($Q_i$) of state *i* was directly derived from the FRET measurement by applying an *E* filter. The resulting fit parameters together with the molecular brightness and triplet state values are reported in Supplementary Table 5.

In a next step, the measurements of $OmpX_{Agg}$ in absence of $Skp_3$ and/or SurA, respectively, were analyzed using the aforementioned parameters. Here we summarized the ensemble of $OmpX_{Agg}$ with different coexisting size in a single diffusion time. While the correlation curve of ~1 µM OmpX without chaperones was fitted with two diffusion components (*k* = 2), three diffusion components (*k* = 3) were used in the presence of chaperones. In the case where both chaperones were present at the same time only three components were used due to the quite similar diffusion times of the $Skp_3$ and SurA bound state. For the error calculation a Jackknifing approach was used, where randomly chosen chunks of the photon stream were removed to measure the variance of the extracted fractions.

### Supplementary Results

#### Denatured state of uOmpX in smFRET experiments

The FRET efficiency ($E$) of refolded $OmpX_{1,149}$ (see Methods) in the presence of 10 mM LDAO under native conditions showed a narrow peak at a high FRET efficiency $\hat{E} \approx 0.9$ (Supplementary Fig. 1a), corresponding to a single conformation, which we attribute to a FRET efficiency of the folded state of $OmpX_{1,149}$ with both dyes in close proximity. After refolding of labelled OmpX in LDAO, we subjected the protein to 6 M GdmCl at 10 pM final concentration. We observed a bimodal distribution in the FRET efficiency histogram (Supplementary Fig. 1b). We attribute the pronounced peak at low FRET efficiency ($\hat{E} \approx 0.2$) to the unfolded conformation of OmpX (uOmpX). The small but distinct second peak at $\hat{E} \approx 0.85$ suggests a compact conformation, different than the folded state. We attribute this state to a misfolded state, since an OmpX aggregation complex is unlikely due to the very low concentration of 10 pM of unfolded $OmpX_{1,149}$ in solution and 6 M GdmCl.

#### Expansion of the $uOmpX_{aq}$ chain on the timescale of hours

The $uOmpX_{aq}$ population changed during the 6 h time course of the experiment and overall shifted towards lower apparent FRET efficiencies ($\hat{E}_{aq}*$) for all the temperatures, thus adapting a more expanded structure. We quantified this expansion using probability distribution analysis (PDA) of the FRET efficiency populations (Supplementary Fig. 1d-f), and acquired $\hat{E}_{aq}*$ as shown in Supplementary Fig. 1g,h (Supplementary Table 2). Such a trend was distinctly perceptible for the measurement conducted at 25 °C, here, the unfolded state shifted from mean FRET efficiencies $\hat{E}_{aq}* \approx 0.62$ at early time points (0–2 h) to $\hat{E}_{aq}* \approx 0.55$ at late time points (4–6 h) (Supplementary Fig. 1g,h). We also observed a broadening of the underlying distance distribution over time from $\sigma_R \approx 0.8$ nm to $\sigma_R \approx 1.0$ nm (Supplementary Fig. 1g), indicating an increased heterogeneity of the $uOmpX_{aq}$ state along with an expansion on a timescale of hours.
